# The nuclear-cytoplasmic ratio controls the cell cycle period in compartmentalized frog egg extract

**DOI:** 10.1101/2024.07.28.605512

**Authors:** Liliana Piñeros, Nikita Frolov, Daniel Ruiz-Reynés, Aleyde Van Eynde, Gabriel Cavin-Meza, Rebecca Heald, Lendert Gelens

## Abstract

Each proliferating cell replicates its DNA and internal components before distributing this material evenly to its daughters. Although the regulation of cyclin-dependent kinases (Cdks) that dictate orderly cell cycle progression is well characterized, how the subcellular localization of the cell cycle machinery contributes to timing is not well understood. We investigated the influence of the nucleus by reconstituting cell cycle oscillations in droplets of frog egg extract in the absence or presence of a nuclear compartment and monitoring dynamics by time-lapse microscopy. We found that the cell cycle time increased in the presence of nuclei, which grew larger with each cell cycle. The correlation between increasing nuclear volume and a longer cell cycle period was maintained across extracts and nuclei from various *Xenopus* species and persisted upon inhibition of DNA replication or transcription. However, inhibition of nuclear import or the kinase Wee1 impacted the relationship between the nuclear-cytoplasmic ratio and the cell cycle period. To conceptually capture these experimental observations, we developed a computational model that incorporates cell cycle oscillations, nuclear-cytoplasmic compartmentalization, and periodic nuclear envelope breakdown and reformation. Altogether, our results support the major role of the nuclear compartment in setting the pace of the cell cycle and provide an explanation for the increase in cell cycle length observed at the midblastula transition when cells become smaller and the nuclear-cytoplasmic ratio increases.

## Introduction

The cell division cycle necessary for the propagation of all species is a highly coordinated series of events that includes replication of DNA and synthesis of cellular components during interphase, followed by an accurate distribution of a complete genome to daughter cells during mitosis (1). At the core of the complex regulatory machinery is the cyclin and cyclin-dependent kinase (Cdk) complex (2) that oscillates in activity for the orderly progression of the cell cycle by phosphorylating hundreds of proteins (3, 4). Counteracting phosphatases, including protein phosphatase 2A and 1 (PP2A and PP1) dephosphorylate Cdk substrates (5–8), generating a complex and dynamic oscillatory interplay that coordinates cell cycle events (9).

Investigation of the cell cycles of the frog *Xenopus laevis* in vivo and in vitro using egg extracts has provided significant insight into the underlying regulatory network (10–18). Rapid cleavage divisions of the early embryo in the absence of cell growth lead to an exponential decrease in cell size with alternating rounds of DNA synthesis and mitosis with periods of about 30 minutes. The core oscillator consists of a negative feedback loop as cyclin B synthesis during interphase reaches a threshold level that activates Cdk1 and mitotic events, which then activates the anaphase-promoting complex (APC/C) that causes cyclin degradation and return to interphase. Additional regulation by various kinases and phosphatases through interconnected positive feedback and double-negative feedback loops introduce nonlinearity phenomena such as ultrasensitivity or bistability in their response functions (19–21). Consequently, the cell cycle oscillator exhibits increased robustness to noise and changes in system parameters (22, 23).

However, the cell encompasses more than just the cytoplasm; subcellular structures also undergo dynamic oscillations during the cell cycle, such as nuclear envelope breakdown and mitotic spindle assembly that have been recapitulated in *Xenopus* egg extracts (24–26). Most studies have focused on a specific event or cell cycle stage, either interphase or mitosis. In this work, we explore how cell cycle duration is influenced by the nucleus, which has been proposed to enhance the robustness of cell cycle oscillations (27, 28) and act as a pacemaker to spatially coordinate the cell cycle via mitotic waves (29–31). Translocation of Cdk1-cyclinB complexes from the cytoplasm to the nucleus introduces resilience to the process of mitotic entry (32) and influences the period of cell cycle oscillations (28). To gain further insight, we utilized cycling frog egg extracts to explore the impact of nuclear size, cell size, and DNA content on the pace of the cell cycle. Our findings reveal a robust correlation between the cell cycle period and the nuclear-cytoplasmic (N/C) volume ratio, even in the absence of DNA replication and gene transcription, challenging the notion that DNA/cytoplasm ratio and/or activation of gene transcription are primary drivers of the lengthened cell cycles at the midblastula transition (MBT). A computational model incorporating nuclear import-export processes and nuclear envelope breakdown conceptually captured the slowing of the cell cycle with increasing nuclear size, in line with our experimental observations. Experimentally perturbing core cell cycle regulators and nuclear transport processes further modulated the scaling relationship between cycle duration and nuclear size.

## Results

### The nucleus slows down the cell cycle in droplets of cycling egg extract

To characterize the impact of the nucleus on cell cycle duration, we reconstituted the cell cycle *in vitro* by encapsulating *Xenopus* egg extracts in droplets of different sizes (33), creating an array of “artificial” cells that were monitored using time-lapse fluorescence microscopy (Fig 1A-B). This droplet setup has been shown to undergo multiple robust cell cycles *in vitro* in the absence or the presence of sperm-derived chromatin (SC) (29, 30, 34). We analyzed individual droplets ranging in diameter from ∼70 to 300 *µm*, mimicking cell sizes observed during the cleavage divisions of early *X. laevis* development. Larger droplets of extract supplemented with sperm chromatin usually contained multiple cell-like compartments with a typical distance between nuclei of about 100 *µm* driven by the microtubules polymerization (24) (Fig. S1; Mov. 1). We also altered the concentration of demembranated sperm chromatin added to manipulate the size of the nuclear compartment within each droplet.

**Fig. 1:**
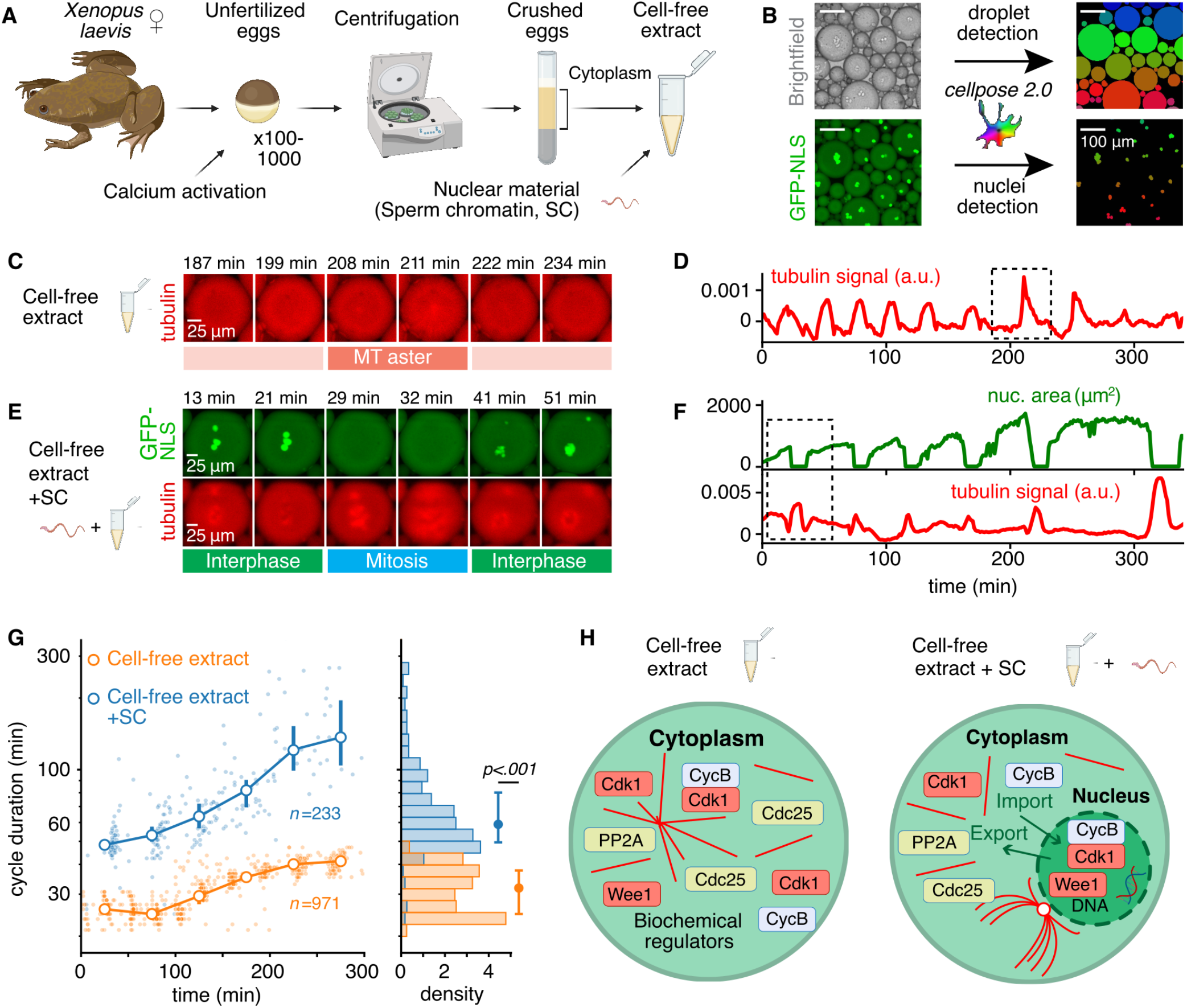
The nuclear compartment slows down the cycle duration in droplets of cycling extract. **A**. Experimental scheme showing the preparation of cycling *Xenopus* extract supplemented with different combinations of sperm chromatin (SC), fluorescently labeled (recombinant) proteins, and drugs. **B**. Example of the droplets and nuclei detection from the acquired brightfield and GFP-NLS images using Cellpose 2.0 (88). Individual detected objects are color-coded with a rainbow palette. **C**. Micrographs of a recurrent microtubule (MT) self-organization visualized by fluorescently labeled tubulin in cycling egg extract without nuclear material. **D**. Respective time trace of the fluorescence intensity of the labeled tubulin in arbitrary units (a.u). **E**. Micrographs of a cell cycle in egg extract with added demembranated SC consisting of periodically alternating phases of presence (interphase) and absence (mitotic phase) of the nuclei shown in bright green (visualized by GFP-NLS). **F**. Respective time traces of nuclear area (green, top) and fluorescence intensity of the labeled tubulin (red, bottom). **G**. Semi-log plot of temporal variation of the cycle duration in cycling extract after adding demembranated SC. Mean *±* s.d. cycle duration in droplets with and without nuclei are shown by hollow circles with error bars. Histograms to the right show the distribution of the aggregate cycle duration for both conditions. Solid circles and error bars indicate the median and interquartile range, respectively. **H**. Model illustrating our hypothesis: nuclear compartmentalization changes the organization, distribution, and dynamics of intracellular content in cell-free extract, thus affecting the pace of the cell cycle.

Applying droplet/nuclei detection algorithms to the acquired microscopy images (Fig. 1B), we first evaluated periodicity in the absence of nuclei by monitoring the intensity of fluorescently-labeled tubulin, reflecting self-organization of the mitotic spindle/aster apparatus (Fig. 1C,D; Fig. S2; Mov. 1), which is driven by the periodic oscillations of cyclin B-Cdk1. During the mitotic phase, robust bundling of microtubules increased the fluorescence signal, which was low during interphase (29, 30, 35). The time trace of the tubulin fluorescent signal in Fig. 1D shows that the cycles of microtubule self-organization represent a regular sequence of prominent peaks with an initial period of around 25 minutes, consistent with the measured cell cycle duration in cleaving *X. laevis* embryos (36–38). Note that starting from around 200 min, peaks in the tubulin intensity signals become sharper, which is associated with the aster formation resembling mitotic spindles (Fig. 1C-E).

We next analyzed the cycling behavior in droplets of *Xenopus* extract supplemented with sperm chromatin and visualized nuclear formation by adding green fluorescent protein fused to a nuclear localization sequence (GFP-NLS), which revealed the periodic alternation of two temporally distinct phases: interphase when the nuclear compartment formed and mitosis when the GFP-NLS appeared diffuse and microtubule fluorescence peaked (Fig. 1E,F; Fig. S2; Mov. 2). We analyzed time traces from one representative extract experiment with 59 droplets containing nuclei and 125 droplets without nuclei. In these droplets, we measured the duration of 233 cycles in droplets with nuclei and 971 cycles in droplets without nuclei. We observed that the cycle duration increased in the droplets containing nuclei, from roughly 50 to 150 min (median 58.94 min). In contrast, in the absence of nuclei, the cell cycle slowed down much less over time (Fig. S3), with the duration starting from 25 min and leveling off at around 40 min (median 31.83 min). This behavior was consistently observed across multiple experimental replicates (See Fig. S4).

These observations suggest a crucial role for the nucleus in slowing down the cell cycle. Based on the existing experimental findings (28–30) and theoretical models (27), we hypothesized that changes in cell cycling behavior are due to partitioning of cell cycle regulators into nuclear and cytoplasmic populations (28, 32, 39), since nuclear import as well as microtubule-driven transport of cytoplasmic regulators towards the nucleus act to increase the density of regulators in the nucleus as it grows (40–43) (right panel in Fig. 1H).

### Cell cycle duration positively correlates with the nuclear-cytoplasmic ratio

If partitioning of cell cycle regulators in the nuclear compartment modulates cell cycle timing, we expected that an increase in N/C volume ratio would slow oscillations, as occurs at the MBT at the onset of zygotic genome activation (37, 44, 45). Strictly speaking, we use the N/C ratio to represent the nuclear-to-cell volume ratio rather than the nuclear-to-cytoplasmic volume ratio; however, we retain this notation as it is commonly used in the field. Note that we used geometry approximations to convert detected droplets/nuclei areas into respective volumes (see Methods and Fig. S5).

To test this hypothesis, we altered the N/C volume ratio by varying the added sperm chromatin concentration (Mov. 2). We observed an immediate formation of larger nuclei in cycle 1 in the extract supplemented with a higher concentration of sperm chromatin (Fig. 2A). Representative time traces of nuclear growth in Fig. 2B show less sustained and slower cycling in extracts with a higher sperm chromatin concentration (See detailed statistics in Fig. S6). The association between nuclear growth and slower cycling is present in all time traces. To test this association, we pooled 376 cycles in droplets containing the same extract but different nuclear densities. We observed a significant positive correlation between the logarithms of cycle duration and N/C volume ratio (Fig. 2C, see legend for statistics). A consistent increase in cycle duration was found with both increased nuclear density and cycle number (Fig. 2D,E). This correlation implies that cycle duration scales as a power of N/C volume ratio. Linear regression confirmed the N/C volume ratio as a key cell cycle duration predictor (*R*^2^ = 0.44, *F* (1, 374) = 289.1, *p* = 1.91e − 48) and revealed the power-law coefficient *– α* = 0.21 *±* 0.012(SE) for the representative experiment in Fig. 2C. Our analysis of various biological replicates revealed that the power-law coefficient (fit slope) could vary between 0.097 and 0.340 (See Table 1 and Fig. 7C). This variation may reflect factors such as egg clutch quality, demonstrating how biological variability can impact cycle dynamics. Examining the effect of the N/C volume ratio across different cycles, we observed that the cycle duration was likely determined by the N/C volume ratio rather than the number of cycles since the start of the experiment (See Fig. S6).

**Table 1:** Summary of the correlation and regression analyses evaluating the relationship between the logarithms of cycle duration and N/C volume ratio in the Control, Interspecies, and Treatment experiments.

| Name | Sample size | Correlation |  |  | Regression |  |  |
| --- | --- | --- | --- | --- | --- | --- | --- |
| | | Pearson's $r$ | $p$-value | $R^2$ | slope | | |
| | | | | | mean±s.e. | 2.5% | 97.5% |
| Control |  |  |  |  |  |  |  |
| Nuclear density (Fig. 2) | 376 | 0.66 | 1.91e-48 | 0.436 | 0.211±0.012 | 0.187 | 0.235 |
| Interspecies, LE x LSD (Fig. 3) | 94 | 0.60 | 1.25e-10 | 0.364 | 0.288±0.040 | 0.209 | 0.267 |
| Wee1 inh., Control (Fig. 5) | 653 | 0.48 | 1.98e-38 | 0.228 | 0.238±0.017 | 0.205 | 0.272 |
| Preformed nuclei, Control (Fig. 5) | 589 | 0.80 | 7.96e-130 | 0.633 | 0.326±0.010 | 0.306 | 0.346 |
| Nuc. import inh., Control (Fig. 5) | 113 | 0.52 | 2.56e-09 | 0.275 | 0.169±0.026 | 0.117 | 0.221 |
| DNA replication inh., Control (Fig. 6) | 263 | 0.52 | 2.22e-19 | 0.267 | 0.097±0.010 | 0.077 | 0.116 |
| Gene transcription inh., Control (Fig. 6) | 113 | 0.66 | 1.83e-15 | 0.436 | 0.243±0.026 | 0.191 | 0.295 |
| Chk1 inh., Control (Fig. 6) | 220 | 0.45 | 2.99e-12 | 0.201 | 0.119±0.016 | 0.087 | 0.15 |
| Wee1 inh. replicate, Control (Fig. S11) | 192 | 0.60 | 8.01e-20 | 0.355 | 0.263±0.026 | 0.212 | 0.314 |
| Nuc. import inh. replicate, Control (Fig. S11) | 139 | 0.43 | 1.23e-07 | 0.185 | 0.340±0.061 | 0.220 | 0.461 |
| DNA replication & Gene transcription inh. replicate, Control (Fig. S11) | 150 | 0.56 | 9.49e-14 | 0.313 | 0.238±0.029 | 0.181 | 0.295 |
| Chk1 inh. replicate, Control (Fig. S11) | 189 | 0.52 | 2.98e-14 | 0.266 | 0.270±0.033 | 0.206 | 0.335 |
| Interspecies experiments |  |  |  |  |  |  |  |
| LE x LEC (Fig. 3) | 49 | 0.72 | 6.87e-09 | 0.514 | 0.257±0.036 | 0.184 | 0.331 |
| LE x TSC (Fig. 3) | 51 | 0.75 | 1.95e-10 | 0.566 | 0.309±0.039 | 0.231 | 0.387 |
| LE x TEC (Fig. 3) | 44 | 0.62 | 6.90e-06 | 0.386 | 0.255±0.050 | 0.155 | 0.355 |
| LE x BSC (Fig. 3) | 45 | 0.70 | 8.31e-08 | 0.491 | 0.326±0.051 | 0.224 | 0.429 |
| LE x BEC (Fig. 3) | 80 | 0.61 | 1.45e-09 | 0.376 | 0.213±0.031 | 0.151 | 0.274 |
| BE x LSC (Fig. 3) | 77 | 0.29 | 9.51e-03 | 0.086 | 0.130±0.049 | 0.033 | 0.227 |
| BE x TSC (Fig. 3) | 108 | 0.50 | 4.08e-08 | 0.248 | 0.184±0.031 | 0.122 | 0.245 |
| BE x BSC (Fig. 3) | 79 | 0.42 | 1.39e-04 | 0.173 | 0.213±0.053 | 0.107 | 0.319 |
| Treatment experiments |  |  |  |  |  |  |  |
| aphidicolin, 2.5 μM (Fig. 6) | 128 | 0.42 | 8.65e-07 | 0.175 | 0.123±0.024 | 0.076 | 0.171 |
| aphidicolin, 2.5 μM, replicate (Fig. S7) | 252 | 0.41 | 2.16e-11 | 0.164 | 0.166±0.024 | 0.119 | 0.213 |
| α-amanitin, 5μM (Fig. 6) | 584 | 0.71 | 3.58e-92 | 0.510 | 0.258±0.010 | 0.238 | 0.279 |
| α-amanitin, 5μM, replicate (Fig. S11) | 605 | 0.56 | 2.10e-50 | 0.309 | 0.225±0.014 | 0.198 | 0.252 |
| SAR-020106, 13nM (Fig. 6) | 119 | 0.42 | 2.53e-06 | 0.173 | 0.166±0.034 | 0.099 | 0.232 |
| SAR-020106, 1μM (not shown) | 38 | 0.59 | 1.04e-04 | 0.346 | 0.189±0.043 | 0.101 | 0.277 |
| SAR-020106, 13nM, replicate (Fig. S11) | 151 | 0.30 | 1.52e-04 | 0.092 | 0.206±0.053 | 0.101 | 0.310 |
| Importazole, 5μM (not shown) | 79 | 0.37 | 8.89e-04 | 0.134 | 0.130±0.038 | 0.104 | 0.205 |
| Importazole, 20μM (Fig. 5) | 71 | 0.035 | 0.772 | 0.001 | 0.008±0.026 | -0.045 | 0.060 |
| Importazole, 20μM, replicate (Fig. S11) | 54 | -0.001 | 0.994 | 0.00 | 0.00±0.051 | -0.103 | 0.103 |
| PD166282, 5μM (Fig. 5) | 50 | 0.06 | 0.680 | 0.004 | 0.007±0.018 | -0.028 | 0.043 |
| PD166282, 5μM, replicate (Fig. S11) | 62 | -0.066 | 0.610 | 0.004 | -0.014±0.027 | -0.069 | 0.041 |
| Preformed nuclei (Fig. S12) | 145 | 0.78 | 2.82e-030 | 0.600 | 0.269±0.018 | 0.233 | 0.306 |
| Preformed nuclei+PD166282, 5μM (Fig. 5) | 302 | 0.72 | 4.12e-049 | 0.515 | 0.149±0.008 | 0.132 | 0.165 |

**Fig. 2:**
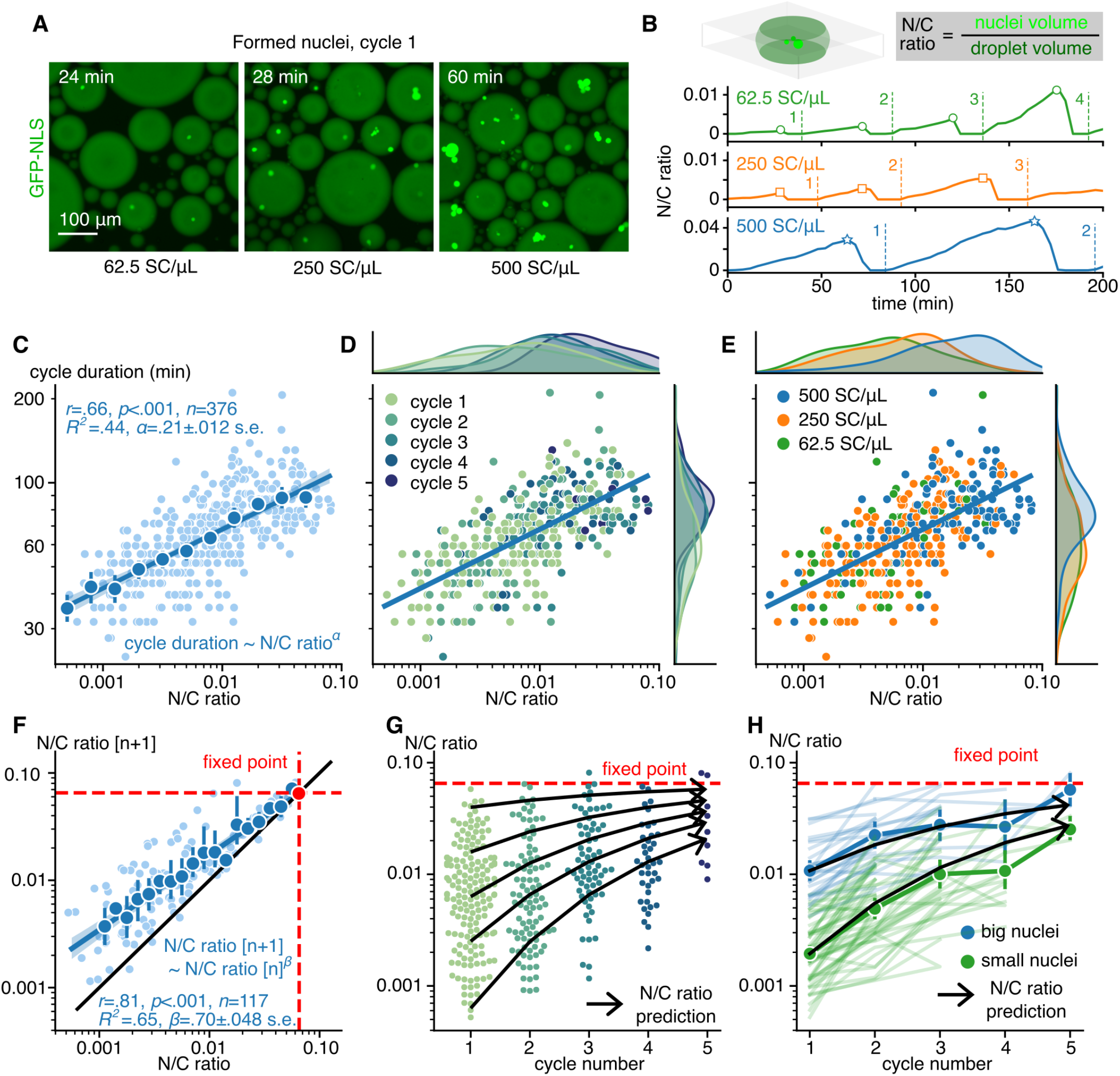
Cycle duration positively correlates with nuclear-cytoplasmic volume ratio. **A**. Cycling extracts were supplemented with different amounts of demembranated sperm chromatin (SC): 62.5, 250, and 500 SC*/µ*L. Micrographs imaging GFP-NLS show the nuclei formed in the droplets of cell-free extracts in the first cycle under different amounts of SC. **B**. Definition of the nuclear-cytoplasmic (N/C) volume ratio (top) and the time traces of nuclear growth in cycling extract with different amounts of SC (bottom). Each cycle is characterized by its duration and N/C volume ratio (indicated with hollow markers). **C**.-**E**. Log-log plots of the cycle duration vs. N/C volume ratio. Blue hollow circles in **C** show means obtained by binning. Error bars are the 95% CIs. The blue line shows the fitted linear model, and shading is the 95% CI of the fit. Pearson’s correlation test was performed, and the goodness-of-fit was assessed. Data in log-log plots **D**-**E** is color-coded with cycle number and SC density, respectively. **F**. Relationship between the N/C ratio at the current *n*^*th*^ cycle and the following (*n* + 1)^*th*^ cycle on a log-log plane. The blue line shows the fitted linear model and shading is the 95% CI of the fit. The black line represents the identity line N/C ratio [*n* + 1] = N/C ratio [*n*]. The red circle at the intersection of black and blue lines indicates a limit of N/C ratio that can be reached within a droplet as predicted by the fitted model. Pearson’s correlation test was performed, and the goodness-of-fit was assessed. **G**.-**H**. Predicting nuclear growth in the droplets using the fitted model from **F**., shown with the bold black arrows. **G**. N/C ratio prediction overlayed with a swarm plot of the experimental data. **H**. N/C ratio prediction overlayed with experimental data trajectories, grouped by the initial size of the nuclei: small (N/C ratio*<* 10^− 2.5^, green) and big (N/C ratio*>* 10^− 2.5^, blue). Thin and bold colored lines represent individual and mean*±*s.d. trajectories in different groups.

By analyzing nuclear size changes across cycles, we revealed that the logarithms of the N/C ratio in the current *n*^*th*^ cycle and the following (*n* + 1)^*th*^ cycle are significantly positively correlated (Fig. 2F, see legend for statistics), providing a simple power-law model for nuclear growth prediction (*R*^2^ = 0.65, *F* (1, 115) = 216.0, *p* = 3.64e − 28). This model aligns with experimental data (Fig. 2G, H), indicating that smaller nuclei grow faster and, strikingly, predicting the upper limit of N/C volume ratio expressed as 10^*γ* / (1*− β*^) where *γ* and *β* are the slope and intercept of the linear regression fit. For the representative experiment in Fig. 2, it approaches 0.065 after multiple cycles (Fig. 2F-H). Combined with the cycle duration correlation (Fig. 2C), this also provides a way to predict the time evolution of cell cycle duration.

### The robust correlation between the cell cycle duration and nuclear-cytomplasmic ratio is preserved across nuclear material from various *Xenopus* species

Our experiments highlighted the crucial role of the nucleus in coordinating cell cycle timing. We next explored the extent to which this phenomenon is conserved among different frog species, specifically comparing the African clawed frog, *Xenopus laevis*, with Marsabit clawed frog, *Xenopus borealis*, both of which are allotetraploid, and the diploid Western clawed frog, *Xenopus tropicalis* (46). These species vary in organismal, cellular, and subcellular sizes, with *X. tropicalis* being the smallest, *X. borealis* of intermediate size, and *X. laevis* the largest of the three (47–49). As nuclear material, we used chromatin extracted from sperm and stage 8 embryos from these three species and prepared cycling egg extracts from each species (Fig. 3A). Note that this is the first time a cycling egg extract has been made with *X. borealis*.

**Fig. 3:**
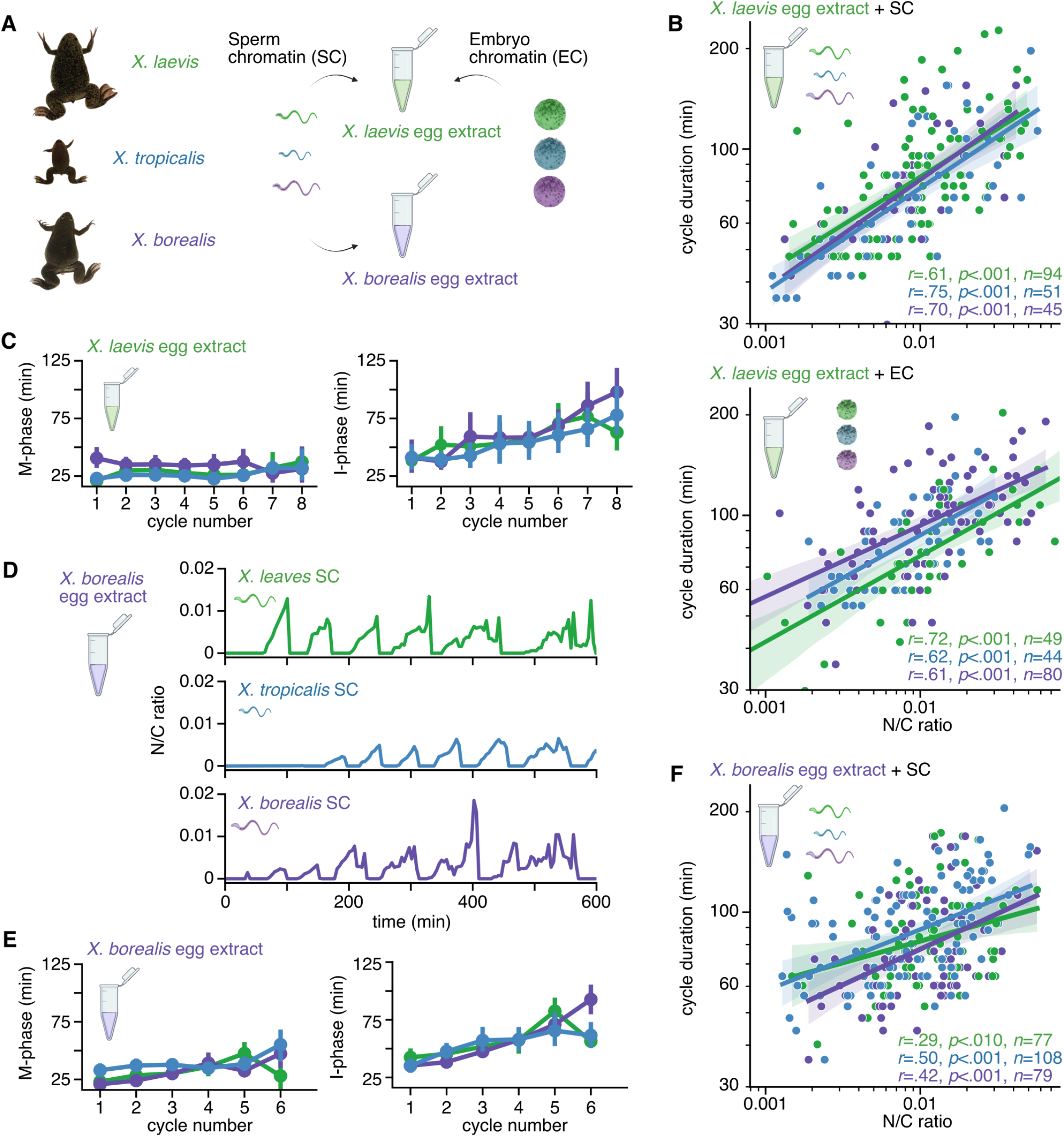
The robust correlation between the cell cycle duration and the nuclear-cytoplasmic volume ratio is preserved across nuclear material from various *Xenopus* species. **A**. Schematic diagram of the cycling extract preparation using *X. laevis* or *X. borealis* eggs, supplemented with one type of nuclear material: sperm chromatin (SC) or stage 8 embryo chromatin (EC, see Material and Methods) from the three frog species: (*X. laevis, X. tropicalis* or *X. borealis*). Frog pictures are adapted from Kitaoka, M., Heald, R., & Gibeaux, R. (2018) (47). **B**. Log-log plot of the cycle duration vs. N/C volume ratio for *X. laevis* cycling egg extract with added SC (top) or EC (bottom) from the three *Xenopus* species. The data is color-coded with the supplemented nuclear material based on the *Xenopus* species: *X. laevis* in green, *X. tropicalis* in blue and *X. borealis* in purple. Pearson’s correlation test was performed, and the goodness-of-fit was assessed. **C**. Duration of the mitotic phase (M-phase, left) and interphase (I-phase, right) vs. cycle number in *X. laevis* cycling extract supplemented with SC and EC. **D**. Time traces of N/C volume ratio in *X. borealis* cycling egg extract with added SC from the three *Xenopus* species mentioned above. **E**. Duration of the mitotic phase (M-phase, left) and interphase (I-phase, right) vs. cycle number in *X. borealis* cycling extract supplemented with SC. **F**. Log-log plot of the cycle duration vs. N/C volume ratio for *X. borealis* cycling egg extract with added SC from the three *Xenopus* species. Pearson’s correlation test was performed, and the goodness-of-fit was assessed.

We observed similar nuclear dynamics in cycling extracts with added chromatin for all three frog species (Mov. 3-5). We first analyzed cell cycle oscillations in *X. laevis* extracts supplemented with either sperm chromatin (Mov. 3) or embryo chromatin (Mov. 4) from the different species (Fig. 3B, Mov. 3). Strikingly, the correlation between cycle duration and N/C volume ratio was found to be preserved across all these different cycling extracts (Fig. 3B, see legend for statistics and Fig. 7C for a comparison across all conditions). As reported *in vivo* (50), the cycle period lengthening in these cycling egg extracts was due to a longer interphase, while the mitotic phase remained of similar duration (Fig. 3C). Similarly, we then analyzed *X. borealis* extracts supplemented with sperm chromatin from the different species (Fig. 3D-F, Mov. 5). Once more, the correlation between cycle duration and N/C volume ratio was maintained (Fig. 3F, see legend for statistics), and the cell cycle was found to slow down due to an increase in interphase duration (Fig. 3E).

While the overall correlation was maintained in all experiments, cell cycle oscillations depended on the type of chromatin added, which determined the size of the nucleus that formed in the first cycle, consistent with Fig. 2E. For instance, sperm chromatin from *X. laevis* formed a larger nucleus that remained similar in size during the first cycles, whereas sperm from

*X. borealis* or *X. tropicalis*, induced the formation of smaller nuclei initially, which grew progressively concomitant with the slowing of the cell cycle period (Fig. 3D). Nuclei formed from embryo chromatin containing maternal and paternal genomic copies were generally larger than nuclei formed from sperm chromatin (Fig S8). For example, diploid *X. tropicalis* displayed more substantial growth (Fig 3B), consistent with earlier findings in *X. laevis* extracts in which smaller nuclei showed faster growth rates (Fig 2H). These results indicated that the relationship between the cycle duration and the N/C volume ratio was preserved across *Xenopus* species. Strikingly, it was even well-reproduced in the inter-species setups and regardless of the source of nuclear material.

### A computational compartment model provides a framework for understanding cell cycle lengthening with in-creasing nuclear size

Our experiments demonstrate that cell cycle duration scales similarly with the N/C ratio independent of initial nuclear density and cycle number. This suggests that the physical volume of the nucleus might drive the cell cycle lengthening directly. We turned to mathematical modeling to examine this hypothesis, explicitly analyzing cyclin B - Cdk1 dynamics using a model by Parra-Rivas et al. (23). The model reproduces periodic oscillations between interphase and mitosis, which are maintained through a balance of negative and positive feedback mechanisms (Fig. 4A). The negative feedback originates from the APC/C’s activation by the cyclin B - Cdk1 complex, subsequently leading to the degradation of cyclin B (51). Conversely, positive feedback is achieved through the regulation of Cdk1 by cell cycle regulatory proteins such as Wee1, Myt1, and Cdc25 (52, 53), which are assumed to be equally distributed in the cytoplasm. These feedback mechanisms not only control Cdk1 activity but also contribute to the activation of APC/C (7, 8, 54), thus establishing multiple intrinsic bistable switches within the regulation of mitosis (21, 23).

**Fig. 4:**
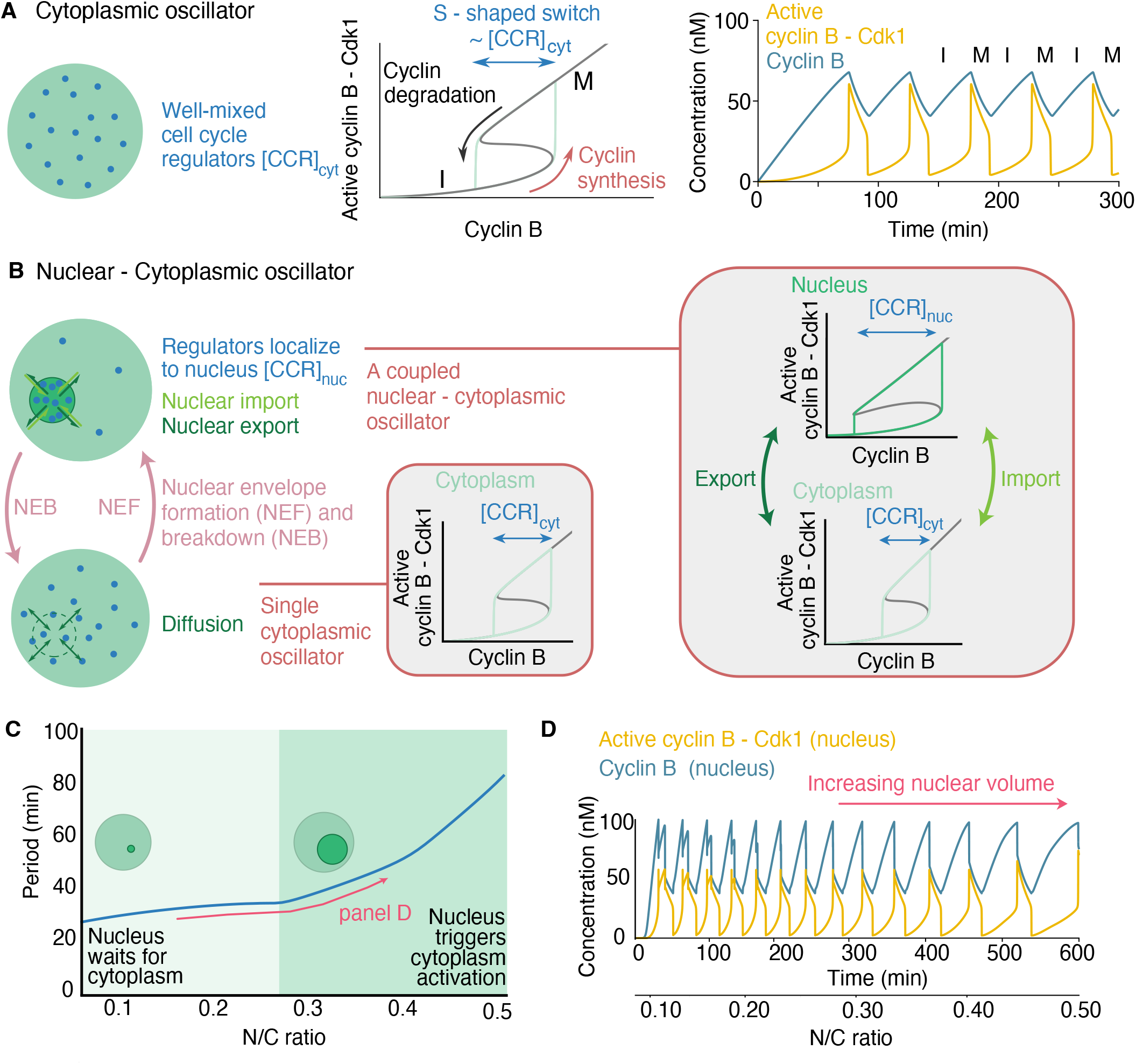
Mathematical modeling shows that nuclear compartmentalization slows down the cell cycle period. **A**. Schematic representation of the cell cycle oscillator describing the periodic accumulation of cyclin B due to synthesis during interphase (I), where the complex cyclin B - Cdk1 rapidly activates as a result of the S-shaped switch and later degraded during mitosis (M). Cell cycle regulator concentration [CCR] controls the S-shaped switch and the relaxation-like dynamics. **B** Compartment model describing the dynamics of two oscillators - the cytoplasm and the nucleus-coupled through import and export during interphase and diffusion during mitosis after NEB. The two compartments exhibit different S-shaped switches due to the compartmentalization of different cell cycle regulators. **C**. Cell cycle period as a function of the N/C ratio, the ratio of the nuclear to cell volumes (blue). Two different regimes are indicated. Small nuclei wait for the cytoplasm to commit to mitosis (light green), while large nuclei trigger the activation of the cytoplasm (green). **D**. Time series for a time-dependent increase in the nuclear volume. The concentration of cyclin B (blue) and active cyclin B - Cdk1 (yellow) in the nucleus is shown.

Nuclear localization significantly impacts cell cycle regulation during interphase, with components like Wee1 residing in the nucleus. Moreover, nuclear envelope breakdown (NEB) and reformation (NEF) also affect the cytoplasmic-nuclear concentration balance. To explore this quantitatively, we adapted the cell cycle model to a two-compartment system representing the cytoplasm and nucleus, linked by cyclin B transport and diffusion post-NEB, forming two interconnected oscillators. Key differences include cyclin B synthesis only in the cytoplasm, a broader bistable switch for the nucleus due to imported cell cycle regulators, and distinct compartment volumes (Fig. 4B).

In this model, constrained by the nuclear import-export rates, cyclin B can accumulate in the nucleus during interphase. Despite requiring a larger cyclin B concentration for Cdk1 activation due to its broader bistable switch, nuclear activation can still occur because of the nucleus’s smaller volume (Fig. S9A). Upon nuclear activation, NEB equalizes the concentrations in both compartments through diffusive mixing, making the oscillatory dynamics in the cytoplasm and nucleus alike (Fig. S9A). Post-NEB, cyclin B degradation continues until the nuclear membrane reforms, completing the cycle (Fig. S9A).

Examining the cell cycle period in relation to nuclear volume, we observe two distinct phases (Fig. 4C). Initially, with a small nucleus, nuclear cyclin B - Cdk1 activates quickly but fails to elevate cytoplasmic levels to the mitotic threshold due to limited volume. As the nucleus enlarges, it shortens the waiting time for cytoplasmic activation, leading to mitosis. Beyond a critical size—about a quarter of the cell’s volume—the nucleus synchronizes cyclin B accumulation rates between compartments, effectively driving the cycle into mitosis by increasing cytoplasmic cyclin levels above the mitotic threshold. This shift also lengthens the cell cycle, influenced by a broader nuclear bistable switch. The model thus explains how a larger nuclear size can extend the cell cycle. It suggests that as the nucleus grows, it shifts the cycle’s control from the cytoplasm to the nucleus, primarily by extending interphase duration (Fig. 4C-D, Fig. S9B-C).

### Targeting regulators of the cell cycle oscillator and nuclear import processes disrupts the observed scaling between nuclear size and cell cycle duration

According to our model’s predictions, we next sought to directly perturb pathways that influence this nuclear-cytoplasmic balance (Fig. 5A) and determine whether these perturbations can disrupt the correlation between N/C volume ratio and cycle duration.

**Fig. 5:**
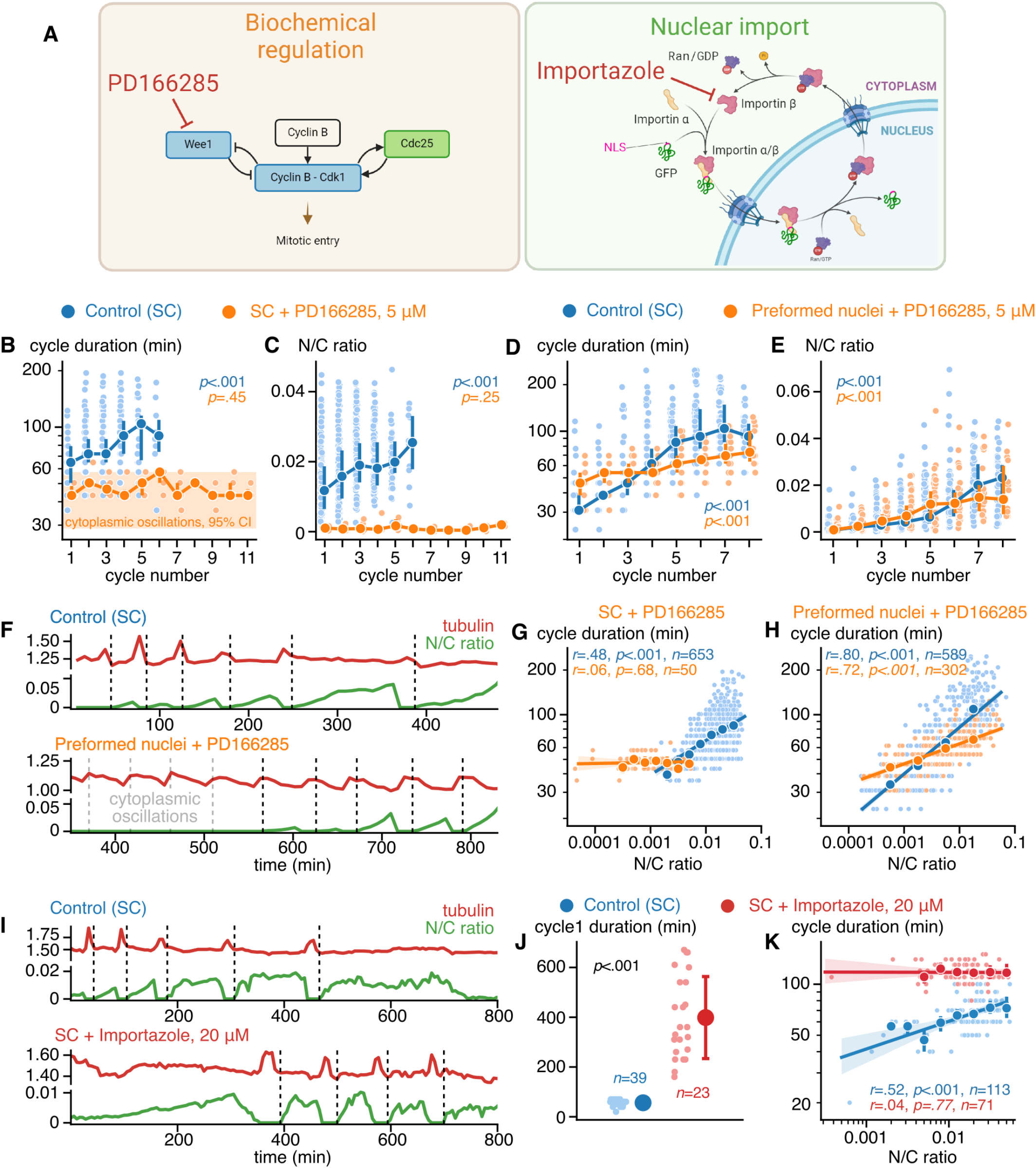
Targeting regulators of the cell cycle oscillator and nuclear import processes disrupt the observed scaling between nuclear-cytoplasmic volume ratio and cycle duration. **A**. Schematic of targeting protein tyrosine kinase Wee1 and nuclear import receptor importin-*—* with respective inhibitors PD166285 and Importazole. **B, C**. Variation of cycle duration and N/C volume ratio against the cycle number in control (SC, *d*_1_=5, *d*_2_=647) and Wee1 inhibition (SC+PD166285, *d*_1_=10, *d*_2_=39) experiments. Orange shading in panel **B** highlights the 95% CI of the cytoplasmic oscillations period, i.e., the cycling period in the droplets of extract without added SC. **D, E**. Variation of cycle duration and N/C volume ratio against the cycle number in control (SC, *d*_1_=7, *d*_2_=582) and Wee1 inhibition with preformed nuclei (Preformed nuclei+PD166285, *d*_1_=7, *d*_2_=295) experiments. In **B-E**, bold lines and dark circles show medians; error bars indicate the 95% CI. The Kruskal-Wallis test was performed. **F**. Typical time series of fluorescent tubulin intensity (red) and N/C volume ratio (green) measured in the cycling extract droplets in control (SC-supplemented) and Wee1 inhibition (SC + PD166285) conditions. Black dashed lines show the start of each cycle; transparent dashed lines show peaking tubulin intensity, indicating cytoplasmic oscillations when the nuclei are not formed. **G, H**. Log-log plots of the cycle duration vs. N/C volume ratio. Bold lines indicate linear regression fit. Dark circles show means obtained by binning; error bars are the 95% CIs. Pearson’s correlation test was performed, and the goodness-of-fit was assessed. **I**. Typical time series of fluorescent tubulin intensity (red) and N/C volume ratio (green) measured in the cycling extract droplets in control (SC-supplemented) and nuclear import inhibition (SC + importazole) conditions. Black dashed lines show the start of each cycle. **J**. Duration of the first cycle in Control and importazole-treated extracts (mean *±* SD). The Mann-Whitney U-test was performed. **K**. Log-log plot of the cycle duration vs. N/C volume ratio. Bold lines indicate linear regression fit. Dark circles show means obtained by binning; error bars are the 95% CIs. Pearson’s correlation test was performed, and the goodness-of-fit was assessed.

First, to investigate the interaction between the nucleus and the proteins in the biochemical network present in both the nucleus and the cytoplasm, the small-molecule compound PD166285 was used to inhibit Wee1 kinase (55, 56), which modulates the width of the bistable switches illustrated in Fig. 4B. Modeling predicts that Wee1 inhibition speeds up cell cycle oscillations. However, in our experimental results without a nuclear compartment (no sperm chromatin added), there was no significant difference in the median cycle period between conditions with (43.22 min) and without (43.22 min) Wee1 inhibitor (Fig. S10A, Mov. 6). This suggests that the period of the cytoplasmic oscillator is not governed by the Wee1-driven bistable switch. One possible reason is that the concentration of cytoplasmic Wee1 may not be sufficient to create a bistable switch that is broad enough to slow down the oscillator’s period. Instead, other mechanisms, such as time delays or alternative bistable switches (23), might play a more dominant role in determining the pace of the cell cycle rather than Wee1.

This behavior may change if a nucleus imports Wee1, leading to a localized increase in its concentration and a broadening of the bistable switch (Fig. 4B). When sperm chromatin was added to the extract, inhibition of Wee1 resulted in a median cycle period of 46.82 min, which was significantly shorter than the median period of 72.04 min in the control condition (Fig. S10A). The dynamics of PD166285-treated extracts supplemented with sperm chromatin were similar to the cytoplasmic oscillator, with less variation in period and a greater number of cycles (*H*(10, 39) = 9.85, *p* = 0.45, Fig. 5B). Moreover, our data indicate a lack of nuclear growth in the droplets treated with the Wee1 inhibitor (*H*(10, 39) = 12.60, *p* = 0.25, Fig. 5C; Fig. S10B). In normal conditions, nuclear and cytoplasmic behaviors are dynamically dependent. When the nucleus is small, the cycling is dictated by the cytoplasmic oscillations, and while it grows, it takes on the leading role from the cytoplasm. Conversely, in the PD166285-treated extract, we propose that fast cytoplasmic oscillations are forced to dominate cellular dynamics, hindering normal nuclear growth. Consequently, the correlation between the cycle period and the N/C ratio is impaired with Wee1 kinase inhibition (Fig. 5G, see legend for statistics). Similar behavior was observed in an independent biological replicate (Fig. S11A).

The underdevelopment of the nuclei under treatment with PD166285, however, questions how functional the nuclei are in these extracts and whether the inhibition of Wee1 consistently disrupts the scaling at larger N/C ratios. To address this, we aimed to pre-form functional nuclei by keeping the extract unperturbed for 40 min, and only after that we treated it with PD166285 (see Methods and Fig. S12A). This manipulation restored nuclear growth in the PD166285-extract to the control level (*H*(7, 295) = 152.72, *p* = 1.09e − 29, Fig. 5E), which was accompanied by a significant but less pronounced cell cycle slowing down (*H*(7, 295) = 127.21, *p* = 2.40e − 24, Fig. 5D). As a consequence, recovering nuclear growth in the Wee1-inhibited extract rescued the scaling between the N/C ratio and cycle duration only partially, with the slope almost twice smaller compared to the control condition (0.149 *±* 0.008 vs 0.326 *±* 0.01, Fig. 5H). Moreover, after treating preformed nuclei extracts with PD166285, nuclei reformed with a significant delay that was preceded by a long phase of rapid cytoplasmic oscillations (Fig. 5F; Fig. S12B, C). It indicates a real-time transition from the cytoplasm-driven oscillations forced by Wee1 inhibition to nuclei-driven cycling facilitated by preformed nuclei. These results, taken together, suggest that Wee1 is crucial but may not be the only cell cycle regulator whose nuclear localization extends cycle duration.

Second, our model suggests that the strength of nuclear import determines the concentration of key regulators in the nucleus and thereby influences the oscillation period. To test this prediction, we added different concentrations of importazole, a chemical nuclear-import inhibitor that targets importin *—*, to the cell-free extract supplemented with sperm chromatin (Fig. 5A,I-K, Mov. 7). While with a smaller dose of the inhibitor (5 *µ*M), the cycle period still increased with a growing N/C ratio, this correlation was completely lost at a higher concentration of 20 *µ*M, although overall cycling periodicity was distinctly extended (Fig. 5K, see legends for statistics). Similar behavior was observed in an independent biological replicate (Fig. S11B). In addition, we observed a significant slowing down of the first cycle in importazole-treated extracts (Fig. 5I, J, see legends for statistics). Considering a global regulatory role of importin *—* in diverse cellular functions (57, 58), its inhibition by importazole can potentially explain systematic cycling slowing down yet the exact molecular mechanism remains unknown. Nevertheless, these results are compatible with our theoretical model’s predictions, underscoring the nuclear compartment’s substantive role in setting cell cycle timing by modulating nuclear and cytoplasmic levels of key cell cycle regulators through nuclear import.

### The cell cycle duration scales with nuclear-cytoplasmic ratio even in the absence of DNA replication, DNA damage checkpoints, and gene transcription

Previous work has shown that the N/C ratio is an important factor in triggering cell-cycle lengthening and initiating gene transcription at the MBT (59–62). Therefore, we aimed to determine if the presence of a nuclear compartment alone can alter the cell cycle period by inhibiting DNA replication, gene transcription, and/or Chk1 activation, which is crucial in the DNA damage response. To do so, we employed specific inhibitors in our experiments with encapsulated egg extract droplets (Fig. 6A, Mov. 8-10).

**Fig. 6:**
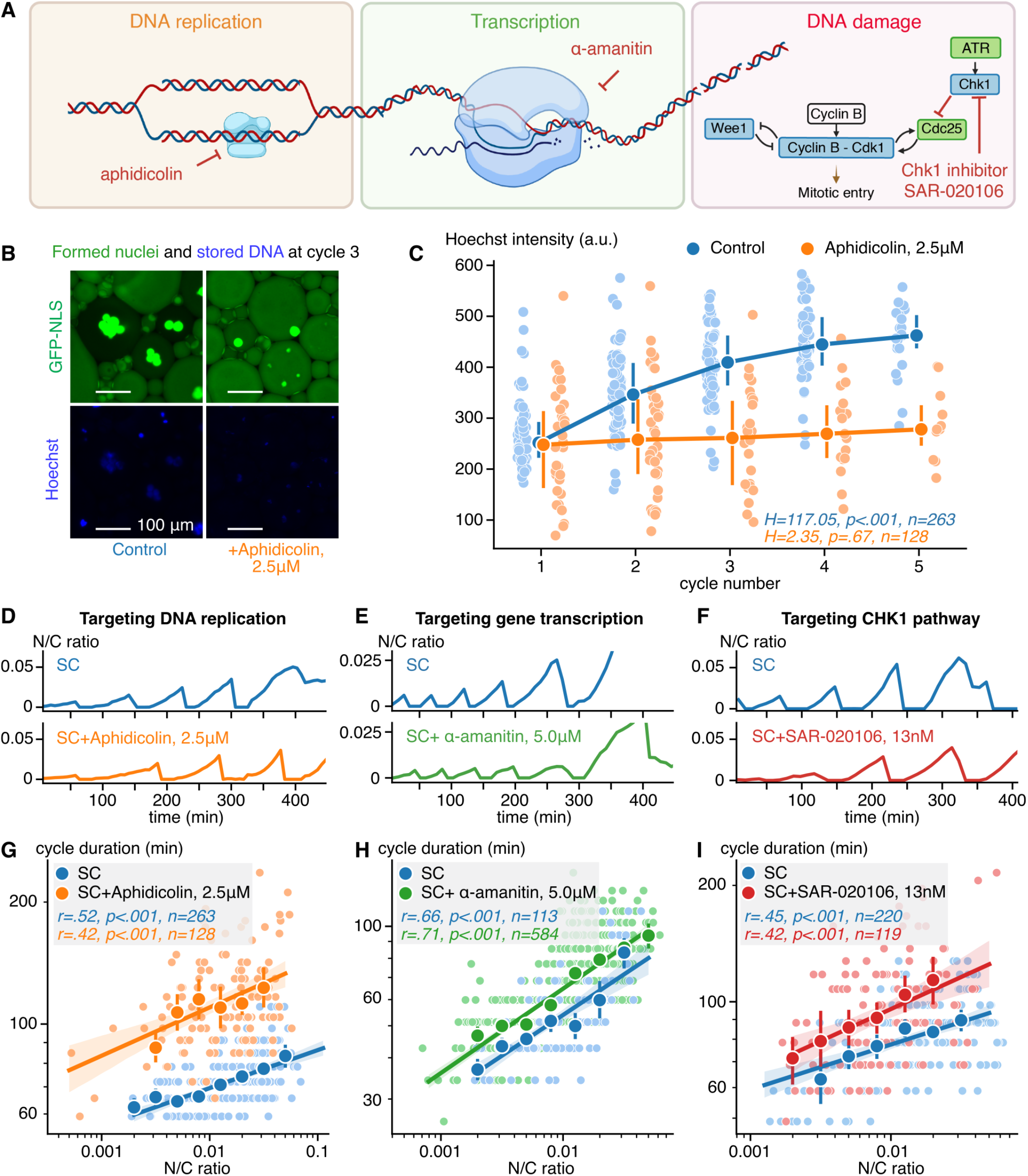
The cycle duration scales with nuclear-cytoplasmic volume ratio in the absence of DNA replication and gene transcription. **A**. Schematic of targeting DNA polymerase, RNA polymerase II/III, and protein kinase Chk1 with respective inhibitors aphidicolin, *–*-amanitin, and SAR-020106. **B**. Micrographs of GFP-NLS (top) and Hoechst (bottom) show the nuclei formed at cycle 3 along with the stored DNA in the control SC extract (left) and extract treated with aphidicolin (right). **C**. Hoechst intensity vs. the cycle number for control SC extract (blue) and extract treated with aphidicolin (orange). Bold lines and dark circles represent medians and error bars show interquartile ranges. The Kruskal-Wallis test was performed. **D-F**. Time traces of N/C volume ratio in control SC extracts (top panels) and extracts treated with aphidicolin, *–*-amanitin, and SAR-020106 (bottom panels in **D-F**, respectively). **G-I**. Log-log plots display the relationship between cycle duration and N/C volume ratio for each treatment: aphidicolin (**G**), *α*-amanitin (**H**), and SAR-020106 (**I**). Dark circles show means obtained by binning. Error bars are the 95% CIs. Lines and shadings represent linear fits and their 95% CIs. Pearson’s correlation test was performed, and the goodness-of-fit was assessed.

**Fig. 7:**
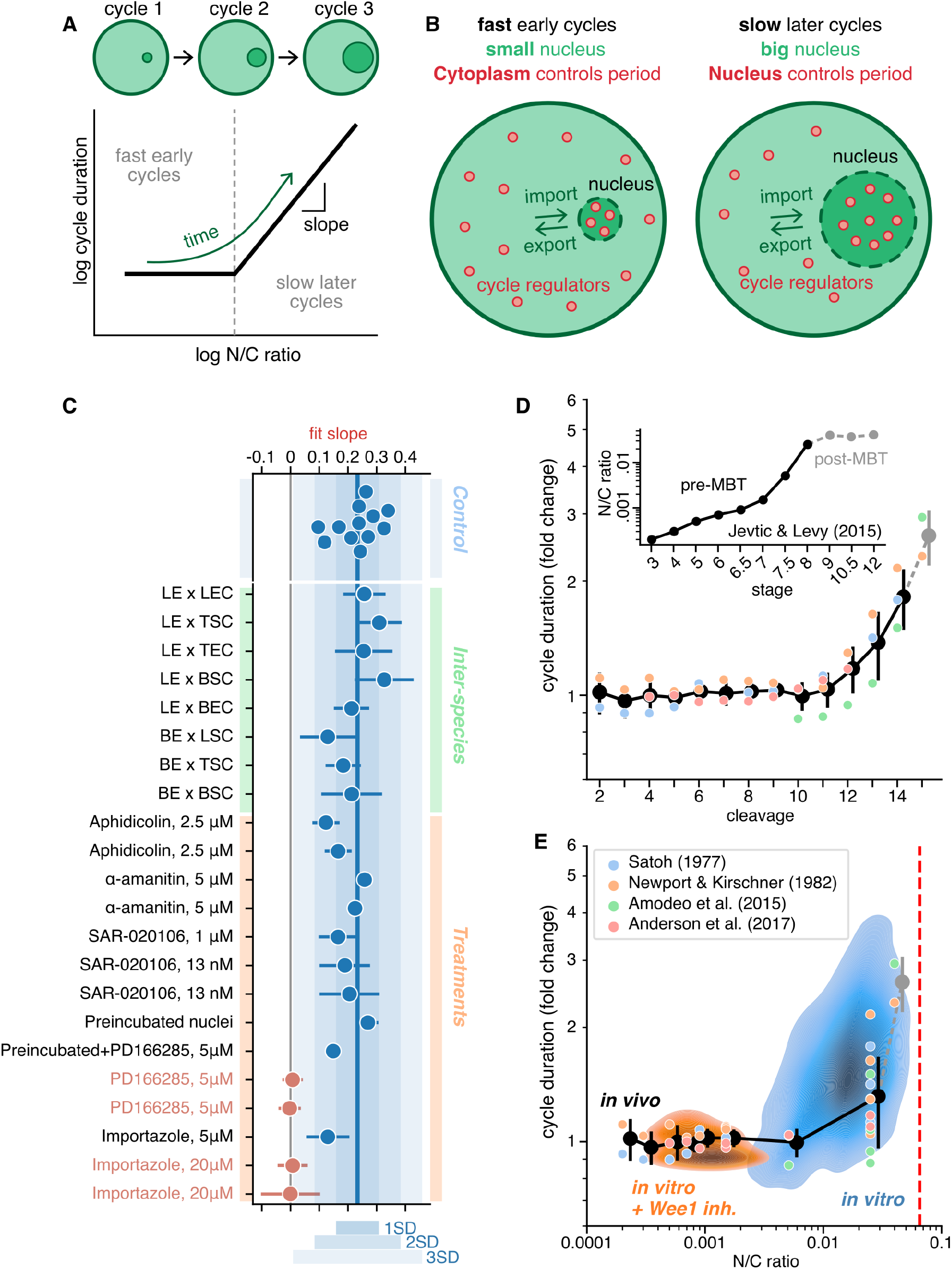
The cell cycle slows close to the midblastula transition due to a combination of i) nuclear compartment-driven changes to the cell cycle oscillator and ii) gene transcription activation. **A**. Cycle duration positively correlates with N/C volume ratio in *X. laevis* cycling egg extracts. **B**. Proposed model of the cycle slowing down due to the relocalization of cell cycle regulators from cytoplasm to nuclei via nuclear import. **C**. Aggregated data of slopes fitted to cell cycle duration vs. N/C volume ratio for all experiments (error bars are the 95% CIs, see Table 1). We collected slopes from 12 control experiments (*X. laevis* extract with SC); 8 slopes from different combinations of *X. laevis* or *X. borealis* extract with SC or EC from *X. laevis, X. borealis*, and *X. tropicalis*; and 14 slopes for *X. laevis* extract with SC with different treatments and their replicates (2.5 *µ*M aphidicolin, 5 *µ*M *α*-amanitin, 13 nM and 1 *µ*M SAR-020106, 5 *µ*M PD0166285, 5 and 20 *µ*M importazole). The blue horizontal line indicates the mean slope across the control experiments; shadings show areas of 1, 2, and 3 respective standard deviations (SD). Two treatments, Wee1 inhibition by PD0166285 and nuclear import inhibition by importazole, considerably decrease or even abolish the correlation between cell cycle duration and N/C volume ratio. **D** Fold change increase in cell cycle duration vs. cleavage stage. Cell cycle period in early *X. laevis* embryos extracted from four sources (36, 38, 59, 67) (see Table 3) and normalized to 35min for the second cell cleavage. The inset shows how the N/C ratio depends on the cleavage stage (45). **E** Cell cycle duration vs. N/C volume ratio for early *X. laevis* embryos (labeled *in vivo*) plotted as a combination of data in panel **D**. Our data obtained using cycling extracts (labeled *in vitro*) is overlayed as density plots: control SC in blue, Wee1 inhibition in orange. The red dashed line represents the predicted steady-state N/C volume ratio for the extract experiments.

Inhibition of DNA synthesis with aphidicolin (63) preserved the level of Hoechst fluorescence intensity with each successive cycle (Fig. 6B-C, see legend for statistics), indicative of DNA replication impairment. However, the consistent correlation between the N/C ratio and cycle duration was nevertheless maintained (Fig. 6D, G, see legend for statistics, Mov. 8).

Next, gene transcription was targeted with *–*-amanitin, a well-known RNA polymerase II/III inhibitor. Blocking gene transcription did not affect cycle lengthening with increasing N/C ratio (Fig. 6E, H, see legend for statistics, Mov. 9), reaffirming its importance in cycle timing, regardless of transcriptional activity.

Lastly, inhibiting the Chk1 pathway to assess DNA damage effects also maintained the correlation between the N/C ratio and cycle duration (Fig. 6F,H, Mov. 10), underscoring the N/C ratio’s pivotal role in cell cycle regulation, independent of these specific pathways. These results are consistent with the theoretically predicted mechanism of cell cycle regulation by nuclear compartment may operate independently of the well-established factors triggering cell cycle lengthening at the MBT.

## Discussion

In this study, we encapsulated droplets containing cycling frog egg extracts in oil to investigate the relationship between nuclear size and the cell cycle. Our findings uncovered a compelling correlation: larger nuclei correlate with a slower cell cycle. Specifically, we identified a robust scaling relationship between the cell cycle period and the N/C ratio (Fig. 7A). Within each extract-containing droplet, we consistently observed nuclei enlarging with each successive cell cycle. Concurrently, the duration of each subsequent cycle increased (Fig.7A). To explain these observations, we proposed a nuclear-cytoplasmic compartment model, according to which nuclear import of cell cycle regulators into a growing nucleus may contribute to slowing the cell cycle (Fig. 7B).

Cell cycle lengthening is well established to occur just before gastrulation at a major embryonic transition known as the midblastula transition (MBT). The MBT is a critical event during early embryonic development in certain organisms, particularly in embryos that undergo cleavage divisions. It marks the transition from rapid, synchronous, reductive, and predominantly transcriptionally silent cleavages to a period of slower, asynchronous cell divisions and the onset of zygotic transcription (64–66). It has long been recognized that the N/C ratio is an important factor in the timing of cell-cycle lengthening and the initiation of gene transcription at the MBT (59, 61, 62). One recognized mechanism involves the titration of maternally loaded factors as the amount of DNA and number of nuclei increase exponentially and cell size decreases exponentially, thereby increasing the N/C ratio (44, 67, 68).

DNA replication during interphase is another crucial process that can directly impact cell cycle duration. The fidelity of DNA replication is closely monitored by various checkpoints to ensure genome stability and proper progression through the cell cycle phases. However, these checkpoints are largely silenced during the early embryo cleavage divisions of many metazoans, facilitating rapid development. Only at the MBT are DNA damage checkpoints activated. Recent work suggests that the activation of the checkpoint kinase, Chk1, is coupled to the N/C ratio via changes in histone H3 availability (69). Such titration of H3 during early cleavage cycles may regulate Chk1-dependent cell-cycle slowing (69, 70).

Remarkably, however, the scaling relationship between the cell cycle period and the N/C ratio we observed remained intact when DNA replication, the DNA damage checkpoint, and gene transcription were inhibited (Fig. 7C, Table 1). Moreover, we demonstrated that this scaling relationship applies to extracts and added nuclear material prepared from different frog species and did not depend on the initial amounts of added sperm chromatin (Fig. 7C). These observations indicate that this cell cycle lengthening is not due to MBT or DNA damage checkpoint activation. Instead, our data indicate that it is a dynamic consequence of the presence of a nuclear compartment. This interpretation is further supported by experiments where inhibition of Wee1 or nuclear import abolished the increase in cell cycle period with N/C ratio (Fig. 7C), as captured by our computational model.

Our work proposes an alternative pathway for lengthening the cell cycle—one that operates independently of regulation through gene transcription or the DNA damage pathway and can manifest before actual zygotic genome activation occurs. In various species, cell cycle periods increase leading up to the MBT (36, 38, 59, 67, 71, 72), while the N/C ratio increases up to 4% as the cells progressively reduce in size (45) (Fig. 7D and Table 3). Strikingly, our data on the scaling of the cycle period with N/C ratio obtained from extract experiments aligns closely with the reported *in vivo* data (Fig. 7E).

Our findings offer new insights into the intricate interplay between nuclear size and cell cycle dynamics, revealing the existence of different mechanisms of temporal control of the cell cycle at various developmental stages. Furthermore, our work could shed light on the connection between the conserved N/C ratio and diseases or other pathological physiological processes such as aging and cancer. Based on the observation that the presence of a nuclear compartment alters the cell cycle period, having very high or very low N/C ratios in cells may contribute to the deregulation of cell cycle progression, potentially associated with altered cellular states involving aging and cancer (73). When cells increase in size, and the surface area-to-volume ratio decreases, their metabolic rate and regenerative capacity diminish, triggering senescence (74–78). Conversely, in cancer, there is uncontrolled cell proliferation, and nuclear size plays a role in the diagnosis and prognosis of different types of cancer (79– 82). Furthermore, recent work by Biswas *et al*. suggests that cells maintain a constant N/C density ratio, driven by pressure balance and osmotic forces across the nuclear envelope, which could provide a physical basis for the conserved N/C volume ratio observed across species and development (83). Such biophysical coupling between nuclear and cytoplasmic compartments may contribute to the dynamic regulation of the cell cycle and offers an intriguing complementary mechanism to the one we propose here.

## Limitations

The current study provides new insights into the role of the nuclear compartment in regulating cell cycle duration. However, several limitations should be acknowledged. First, our imaging approach was performed in two dimensions by averaging fluorescence intensity over z-stacks, which limits the ability to fully resolve the three-dimensional structure of nuclei and droplets. To estimate volumes from detected areas, we made geometric assumptions — approximating droplets as spheres or cylinders depending on their size — which may introduce additional measurement errors, particularly in larger droplets. These approximations, while necessary, add uncertainty to precise N/C ratio calculations.

Second, the experimental data show considerable variation, which in some cases resulted in overlapping confidence intervals for fitted linear regressions. This variability makes it difficult to confidently conclude differences in scaling relationships between some experimental conditions, such as interspecies comparisons. Therefore, we have refrained from making strong claims about such differences, unless the trends are statistically robust.

Third, while our computational model provides a useful conceptual framework for understanding how nuclear-cytoplasmic compartmentalization could influence cell cycle dynamics, it does not quantitatively reproduce experimental findings. The model was designed to capture the conceptual relationship between increasing N/C ratio and cell cycle period lengthening. However, it assumes constant nuclear import and export rates, independent of nuclear size, and does not account for potential feedback mechanisms such as size-dependent transport rates or spatial regulation of cyclin B dynamics. Within the experimentally observed N/C ratio range (0.01–0.1), the nucleus in our model remains too small to substantially influence the cell cycle period unless unrealistically high nuclear import rates are invoked. Moreover, the volume ratio required in the model for the nucleus to significantly affect the cell cycle is an order of magnitude higher than experimentally observed values. This discrepancy may reflect the fact that our measurements do not account for the functionally available volume, as cellular space is partially occupied by organelles, macromolecules, and chromatin. Consistent with this, estimates of nuclear/cytoplasmic volume ratios using FRAP in yeast are around 0.1, while geometric estimates typically range from 0.01 to 0.1 (with N/C ratios between 0.06 and 0.1) (84). An additional limitation is that our model does not capture spatial phenomena such as mitotic trigger waves. In spatially extended systems, smaller nuclei may locally initiate mitotic activation, which then spreads through the cytoplasm as a wave, a process that our compartment-based model cannot simulate. Furthermore, our model does not incorporate contributions from other known regulators of cell cycle timing, such as other key kinases and phosphatases. Thus, while the model is a valuable conceptual tool, further theoretical refinements and experimental validations are required to fully describe the complex interactions governing nuclear-cytoplasmic regulation of the cell cycle.

Another important limitation is that although our combined experimental and modeling efforts strongly support a role for the nuclear compartment in modulating cycle duration, these observations do not establish strict causality. It remains possible that the observed scaling arises from indirect or correlated processes rather than direct nuclear control. Dissecting these relationships will require future experiments designed to test causality more rigorously.

Finally, specifically impairing nuclear import without off-target effects is technically challenging. Importins, which mediate nuclear import, are also involved in other essential processes such as cell cycle progression. Consequently, inhibition of nuclear import may lead to unintended side effects — such as generalized slowing of the cell cycle — that could obscure the specific impact of disrupting nuclear import on N/C ratio scaling. These limitations highlight areas for further experimental refinement to isolate and verify the direct contributions of nuclear import processes to cell cycle control.

## Methods

### *Xenopus laevis* and *Xenopus borealis* egg extract preparation and nuclear assembly

Frog handling and preparation of cycling *Xenopus laevis* egg extracts were performed as described in Nolet et al., 2020, following the protocol of Murray, 1991 (29, 85). After dejellying, eggs were activated using calcium ionophore A23187 (0.5 µg/mL in 0.2 × Marc’s Modified Ringer’s buffer, Sigma-Aldrich) for 2 min to start the biochemical processes of the cell cycle *in vitro*. The ethical committee approved the frogling, including the one injection and the egg collection, under Project 107-2021.

The extract was supplemented with different concentrations of *Xenopus* demembranated chromatin prepared from sperm or stage 8 embryos from three frog species: *X. laevis, X. tropicalis* and *X. borealis*. The supplemented nuclear material induces spontaneous nuclear formation. Sperm chromatin was prepared according to the protocol by Murray, 1991 (85). Briefly, demembranated sperm chromatin was prepared from isolated sperm by treatment with lysolecithin, which disrupts the plasma membrane of the cells and the nuclear envelope of the nucleus. Similarly, embryo chromatin was prepared from stage 8 embryos according to the protocol by (86) and treated with lysolecithin. To improve optical transparency, the extracts were clarified twice at 16000 g for eight minutes (30). Cytochalasin B (10 µg/mL, Sigma-Aldrich) was added to inhibit actin assembly and thus gelation-contraction, keeping the extract fluid at room temperature (87).

The cycling *Xenopus borealis* egg extracts were prepared utilizing the same protocol. The cycling extract is functional after being kept on and off the ice during incubation. Prior to adding DNA and forming droplets, the *X. borealis* extract was kept at 16°C. In some specific experiments, additional purified proteins, fluorescence-labeled reporters, and drugs were added to the extract to influence and visualize the cell cycle dynamics, as described below.

### Extract supplements

We used two probes to follow the oscillations between interphase and mitosis: green fluorescent protein with a nuclear localization signal (GFP-NLS) and fluorescence-labeled tubulin. The GFP-NLS accumulate in the nuclei formed during interphase in the extract after adding nuclear material like sperm chromatin or pure DNA and disperse after nuclei break down in the mitosis. The microtubules’ fluorescence indicates the labeled tubulin’s polymerization driven by the activity of Cdk1 in the extract, even in the absence of nuclear material. The kymograph shows the oscillation of the fluorescent microtubule signal.

Recombinant GST-GFP-NLS (referred to as GFP-NLS) was bacterially expressed and purified by glutathione agarose. GFP-NLS was added to the extract at ∼25 *µ*M to visualize the oscillations between the interphase and mitotic phase, following the formation of the nuclei. The construct for GFP-NLS was kindly provided by James Ferrell (Stanford Univ., USA). The polymerization dynamics of the microtubules and the DNA were followed using fluorescently labeled porcine tubulin (HiLyte Fluor™ 647; Cytoskeleton, Inc) at 1 *µ*M final concentration and DNA staining (Hoechst 33342) at 5 *µ*g/mL final concentration, respectively.

The following reagents were added to the extract, as indicated in the text, to study the perturbations of the system that could impact the cell cycle period. Aphidicolin was used as an inhibitor of DNA replication at final concentrations between 0.5 and 10 *µ*M. SAR-020106 13 nM was used as a selective Chk1 inhibitor. *α*-amanitin was tested in a range of 2.5 and 10 *µ*M to inhibit the DNA transcription. Wee1 kinase was inhibited by PD0166285 5 *µ*M.

For the experiments with preformed nuclei, the extract was kept at room temperature until the beginning of the interphase on the first cell cycle oscillation. The Wee1 kinase inhibitor PD0166285 5 *µ*M was added after the first nuclear formation. Finally, nuclear import was inhibited by adding importazole (Sigma-Aldrich), an inhibitor of importin *—*, in concentrations of 5, 20, and 40 *µ*M.

### Microscopy data acquisition

We used the setup described in Guan et al., 2018. to obtain water-in-oil microemulsions of *Xenopus* egg extract to characterize the cell cycle dynamics in vitro (33). For the following experiments, the vortexing method was used to generate the droplets of egg extract that allow following their individual oscillations by time-lapse microscopy. The vibration speed and duration (5 s) were adjusted to achieve droplet sizes ranging from 70 *µ*m to 300 *µ*m. For imaging, the droplets were loaded in a Teflon-coated microchamber prepared by capillary effect and placed in a glass-bottom 6-well plate covered by mineral oil. The samples were imaged in time-lapse using a Leica TCS SPE confocal fluorescence microscope (5x objective). The images were recorded in bright-field and multiple fluorescence channels at an acquisition time interval of 1–7 min for 18 hours at room temperature. The microscopy file was saved in `.lif` (Leica Image File) format.

### Image analysis

Acquired fluorescent microscopy videos were analyzed to extract the time-resolved intensity of microscopy reporters staining different intracellular processes from each individual artificial cell (droplet of cell-free extract). Generally speaking, image analysis was divided into three stages – (i) image segmentation, (ii) transformation of images into time series, and (iii) cell cycle quantification. A detailed overview of the image analysis steps is given below.

### Image segmentation and object tracking

Firstly, `.lif`-files containing analyzed microscopy videos were saved as image sequences in separate folders per channel per position using `readlif` package for Python. Secondly, these images were segmented using a Cellpose 2.0 algorithm (88) to detect droplets and respective nuclei. We have chosen this software because, on the one hand, it offers a zoo of built-in models for detecting cellular and subcellular objects. On the other hand, it is flexible enough to annotate new data and train a custom model based on the existing ones. Specifically, to batch analyze acquired image sequences, we used `serialcellpose` – a Cellpose 2.0 implementation for Napari (89, 90). For the segmentation of droplets, we used the brightfield microscopy images and applied a pre-trained cytoplasm model `cyto2`. To segment nuclei, we used the images of the GFP-NLS reporter and trained a custom model starting from a built-in model `nuclei`. Features of the detected objects’ morphology, e.g., roundness, area, equivalent diameter, etc., were automatically produced by the `serialcellpose` plugin and saved in respective `.csv` files (one file per image). To complete image segmentation, we tracked detected objects in each video using the custom code based on the commonly used intersection-over-union metric (IoU). Noteworthy, artificial cells smaller than 3600 *µm*^2^ (or equivalently less than 68 *µm*^2^ in diameter) and having roundness *<* 0.75 were excluded from the dataset.

### Converting detected object areas into volumes

Time-lapse fluorescent microscopy conducted in this study produces sequences of 2D images by averaging over z-stacks; thus, we lack information about the 3D structure of detected droplets and nuclei. To convert measured areas of droplets and nuclei detected in 2D images into respective volumes, we made the following assumptions about objects’ geometry (Fig. S5):

- *Droplets*, whose equivalent diameter was <u>smaller</u> than the glass capillary thickness of <u>100 *µ*m</u>, were approximated as *equivalent spheres*;
- *Droplets*, whose equivalent diameter was <u>larger than</u> the glass capillary thickness of <u>100 *µ*m</u>, were approximated as *equivalent cylinders*;
- *Nuclei* were <u>always</u> approximated as *equivalent spheres* since their equivalent diameter is smaller than the glass capillary thickness of 100 *µ*m.

In the *sphere* approximation, volumes were computed as:

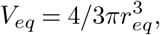

and in the *cylinder* approximation:

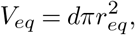

where 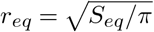 is equivalent radius of a detected object and *d* = 100 *µ*m is the glass capillary thickness.

### Transforming images into time-series

To analyze how the periodicity of cycling cell-free extract depends on the nuclear size, we computed the following time series for each detected and tracked droplet:

- *nuclear-cytoplasmic ratio (N/C ratio) quantifying the nuclear growth*. It is computed by simply dividing the total volume occupied by detected nuclei within the detected droplet by the detected droplet volume:

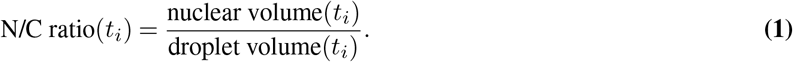
- *intensity of the Hoechst stain within nuclei representing the amount of nuclear DNA*. By analogy with GFP-NLS, it is computed from the Hoechst images as a sum of the brightnesses of each pixel attributed to detected nuclear objects within the droplet:

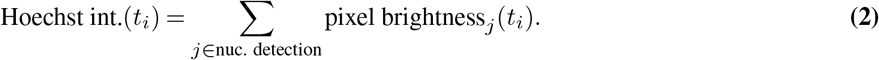
- *intensity of the tubulin reporter*. Active polymerization of microtubules during mitosis induced by the Cdk1 activity is reflected as increased intensity of the tubulin reporter (Fig. 1C-D). During the m-phase, it is usually localized in the center of the cellular compartment, while the brightness of the compartment’s periphery remains primarily unchanged. Assuming that the droplet is a round object and the microtubule polymerization is radially symmetric, we compute averaged tubulin reporter intensity in a radially floating circle of a width *w* ≈ 4*µm* and an inner radius *R*; this results in the “radial” kymograph of tubulin reporter intensity tubulin int.(*R, t*_*i*_). The tubulin signal characterizing the periodicity of cycling extract oscillations in individual droplets is acquired by averaging tubulin reporter intensity in the area of highest polymerization, i.e., within the *R*_0_ ≈ 10*µm* from the droplet center:

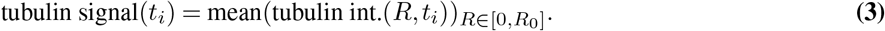

### Cell cycle quantification

From constructed time-series, we can identify the cell cycle events (see example in Fig. 2B) and quantify their duration, size of the formed nuclei, and intensity of the reporters. We introduced the following quantities to characterize the cell cycles and analyze the dynamics of cycling extracts in different experimental conditions:

- *duration* of the *n*^*th*^ cycle (min) was measured from the N/C ratio time-series as the time interval between the starting points of the *n*^*th*^ and the (*n* + 1)^*th*^ interphases, where N/C ratio *>* 0 (Fig. 1F, Fig. 2B). For conditions without injection of nuclear material, cycle duration was measured from the tubulin intensity time series as the time interval between the peaks of intensity reflecting the Cdk1-induced microtubule polymerization (Fig. 1E).
- *interphase duration* of the *n*^*th*^ cycle (min) was measured from the N/C ratio time-series as the time interval in which N/C ratio *>* 0 during this cycle, meaning that the nuclear compartment is formed (Fig. 1B).
- *mitosis duration* of the *n*^*th*^ cycle (min) was measured from the N/C ratio time-series as the time interval in which N/C ratio =0 during this cycle, meaning that the nuclear compartment is absent (Fig. 1B).
- *N/C ratio* of the *n*^*th*^ cycle was measured as a maximal N/C ratio reached during this cycle.
- *Hoechst intensity* of the *n*^*th*^ cycle (a.u.) was measured as a maximal intensity of Hoechst reached during this cycle.

Each of the quantities mentioned above is associated with the respective cycle number *n* = 1, 2, 3, … and the respective starting time of the cycle (min).

### Statistical analysis and modeling

Differences in the samples of collected cell cycle quantifiers between experimental conditions were analyzed using appropriate statistical tests. The normality of the data samples was determined using the Shapiro-Wilk test. Since most of the data did not meet the normality assumption, non-parametric techniques for independent samples — the Mann-Whitney U-test or Kruskal-Wallis test — were used. For each test, we indicate *p*-values, respective statistics values, and degrees of freedom.

To test the linear relationship between two variables of interest, we used Pearson’s correlation coefficient and respective *p*-value. We applied linear regression to model this relationship using the ordinary least squares method. For each fit, we compute the fit’s slope and intercept, their 95% CIs, and the goodness-of-fit *R*^2^.

All statistical tests and linear regressions were performed in Python using the `scipy.stats` and `statsmodels.regression` packages.

### Dynamical model

The model describes the dynamics of the main cell cycle regulators when compartmentalized in the cytoplasm and the nucleus. The model uses in its core the cell cycle model previously introduced by Parra-Rivas et al. (23) describing the periodic oscillations in the concentration of cyclin B (*c*), the active fraction of cyclin B-Cdk1 (*c*1*/c*), and the activity of the anaphase-promoting complex/cyclosome (APC/C, *a*). Here, we adapt it to describe the concentration of the active cyclin B-Cdk1 (*c*1) instead, assuming cyclin B-Cdk1 activates much faster than other contributions, increasing the activity. Thus, the time evolution of *c, c*1, *a* in each compartment is described by the following ODEs:

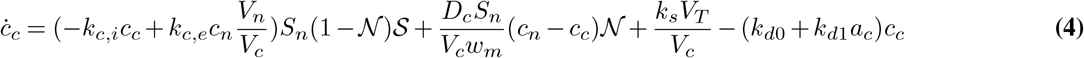

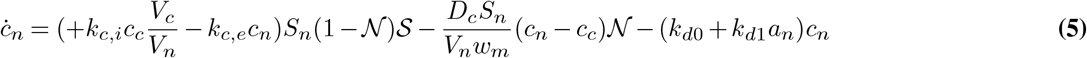

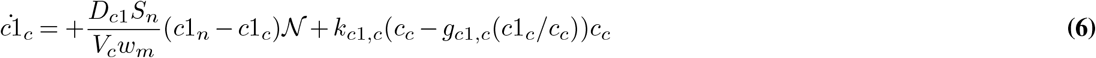

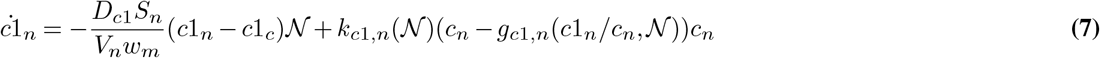

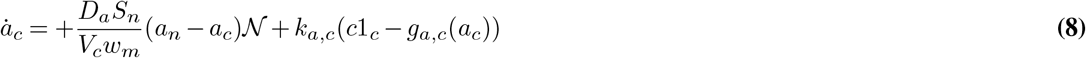

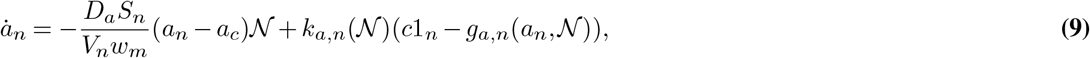

using the subscripts α ≡ {*c, n*}, such as *c*_*c*_ and *c*_*n*_ respectively for cyclin B in the cytoplasm and the nucleus and equivalently for the other variables. Import and export processes have only been considered for cyclin B, which is the main component translocated into the nucleus and occurs at the rate *k*_*c,i*_ and *k*_*c,e*_ accounting for the surface of the nucleus *S*_*n*_, and the different compartment volumes -the cytoplasm volume *V*_*c*_ and the nuclear volume *V*_*n*_-in Eqs. (4-5). The surface of the nuclear volume can be written in terms of the nuclear volume assuming spherical geometry as 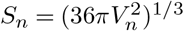. The total volume of the cell

*V*_*T*_ = *V*_*c*_ + *V*_*n*_ is considered constant and estimated from the experimental data as 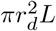, where *r*_*d*_ is the radius of the nucleus and *L* the height of the microchamber. The model accounts for nuclear envelope breakdown with the variable 𝒩 ∈ [0, 1]. When the envelope is broken 𝒩= 1, import-export terms are suppressed, and the diffusive term is active, balancing the concentration in the two compartments. When the nuclear envelope closes 𝒩= 0, the situation reverses with diffusion suppressed and import and export active. The transport through diffusion is controlled by *D*_*β*_, where the subscript refers to each variable *β* ≡ {*c, c*1, *a*}, and the width of the membrane *w*_*m*_, which accounts for the separation between the concentration inside the nucleus *c*_*n*_ and outside *c*_*c*_ leading to a concentration gradient (*c*_*n*_ − *c*_*c*_)*/w*_*m*_. For simplicity, here, the diffusion has been considered equal for the three variables on the typical scale of proteins between 70-250 kDa (91, 92). Nuclear envelope breakdown is triggered when the activity is high either from the inside or outside the nucleus. NEB is described using an ultrasensitive response dependent on the active fraction of cyclin B-Cdk1 in both compartments of the form:

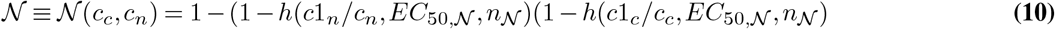

which becomes equal to one when either the active fraction in the cytoplasm *c*1_*c*_*/c*_*c*_ or the nucleus *c*1_*n*_*/c*_*n*_ is greater than *EC*_50,𝒩_ following the Hill response,

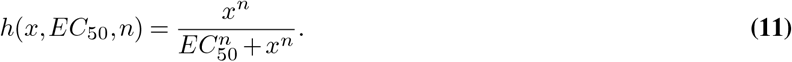

Spatial positive feedback (SPF) regulating import-export terms (32) has been described for simplicity using an ultrasensitive response,

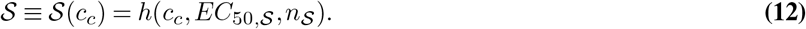

Cyclin B is synthesized only in the cytoplasm at a rate *k*_*s*_ and degraded by APC in both compartments at a rate (*k*_*d*0_ + *k*_*d*0_*aα*). The activation of cyclin B-Cdk1 follows a bistable response curve *g*_*c*1,*α*_ dependent on the active fraction *c*1_*a*_*/cα* described by the cubic expression,

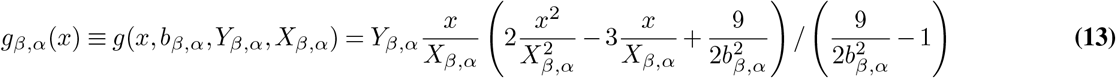

where *b*_*β,α*_ ∈ [0, 2] controls the width of the bistable response curve and *X*_*β,α*_ and *Y*_*β,α*_ are scaling factors for (*x, y* = *g*_*β,α*_ (*x*)) respectively. *g*_*β,α*_ (*x*) is written in a general form to account for different bistable response curves for the different compartments and variables as shown in Eqs. (8-9). Thus, the bistable response curves *g*_*a, α*_ (*x*) describe the APC/C switch in each compartment.

When NEB occurs, the bistable response in the nucleus must become equivalent to the bistable response of the cytoplasm to ensure the nuclear compartment completely vanishes. We describe this effect by introducing the dependence of the bistable response *g*_*β,α*_ (*x*) on the nuclear envelope variable 𝒩, which read:

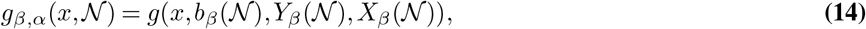

where the dependence of the parameters is given by:

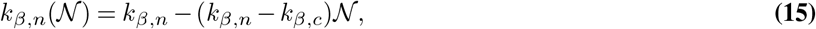

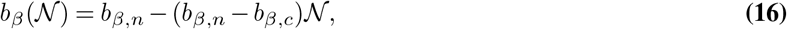

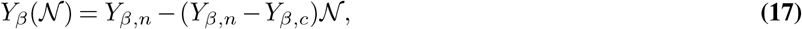

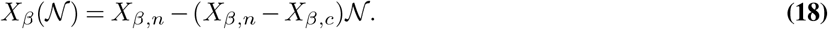

The parameters used for the numerical simulations in this work have been taken from the literature when available and are given in Table 2.

**Table 2:**
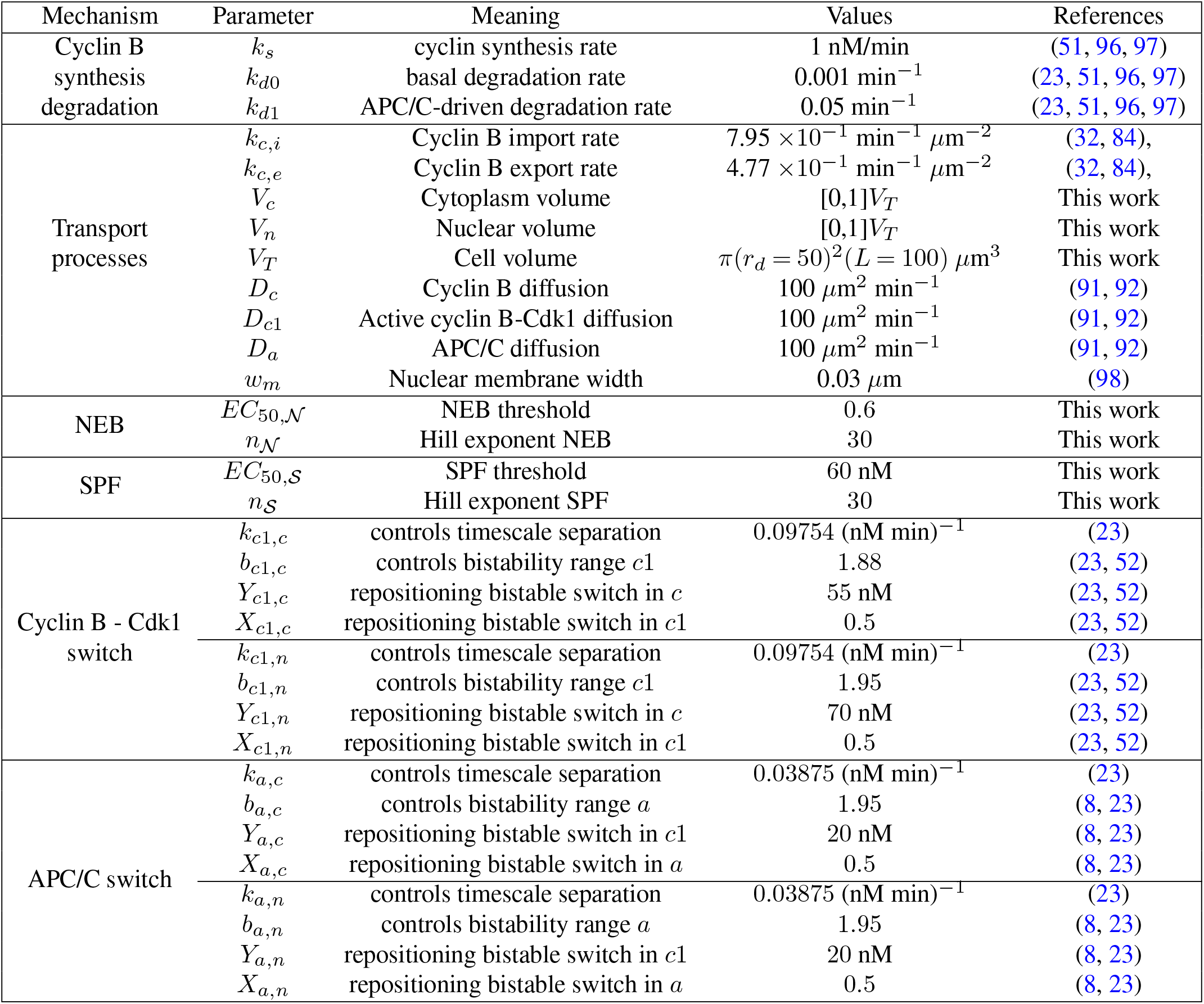
Parameters used in the compartment cell cycle oscillator model (4)-(9).

**Table 3:** Meta-analysis of the *in vivo* studies measuring nuclear-cytoplasmic (N/C) volume ratio and cycle duration at different developmental stages in *X. laevis* eggs.

| cleavage | stage | N/C ratio (%) |  | cycle duration (min) |  |  | mean (min) |
| --- | --- | --- | --- | --- | --- | --- | --- |
| 2 | 3 | 0.02 | 32.5 | 38.85 | - | - | 35.68 |
| 3 | 4 | 0.03 | 31.4 | 36.25 | - | - | 33.83 |
| 4 | 5 | 0.05 | 31.4 | 38.85 | - | 34.69 | 34.98 |
| 5 | 6 | 0.07 | 32.6 | 35.85 | - | 35.05 | 34.5 |
| 6 | 6.5 | 0.09 | 37.6 | 36.25 | - | 33.62 | 35.82 |
| 7 | 7 | 0.15 | 33.9 | 38.45 | - | 33.94 | 35.43 |
| 8 | 7 | 0.15 | 35.6 | 38.15 | - | 33.74 | 35.83 |
| 9 | 7 | 0.15 | 35.9 | 37.75 | - | 34.69 | 36.11 |
| 10 | 7.5 | 0.51 | 36.4 | 36.25 | 30.36 | 36.45 | 34.86 |
| 11 | 8 | 2.5 | 39.4 | 36.65 | 30.76 | 38.36 | 36.29 |
| 12 | 8 | 2.5 | 45.4 | 45.25 | 33 | 41.07 | 41.18 |
| 13 | 8 | 2.5 | 49.6 | 57.25 | 37.66 | - | 48.17 |
| 14 | 8 | 2.5 | 62.4 | 75.65 | 52.68 | - | 63.58 |
| 15 | 9 | 4 | - | 81.25 | 102.92 | - | 92.09 |
| - | Xenbase | Jevtic (93) | Satoh (38) | Newport (60) | Amodeo (67) | Anderson (36) | - |

### Meta-analysis of *in vivo* studies

To compare our *in vitro* results with *in vivo* studies of early embryonic cell cycles in *X. laevis*, we aggregated the dependencies of cell cycle duration on cell cycle/developmental stage for previous works (36–38, 67). From Jevtic and Levy (93), we acquired the association between the N/C ratio and the cell cycle/developmental stage *in vivo*. The collected data are presented in Table 3.

## Data and code availability

- All experimental data and codes have been deposited at Gitlab (94), and are publicly available as of the date of publication.
- All original modeling code has been deposited at Gitlab (95), and is publicly available as of the date of publication.
- Any additional information required to reanalyze the data reported in this paper is available from the lead contact upon request.

## Acknowledgements

L.G. acknowledges funding by the KU Leuven Research Fund (grant number C14/23/130) and the Research-Foundation Flanders (FWO, grant number G074321N). L.P. acknowledges funding by the Research-Foundation Flanders (FWO, personal fellowship - grant number 11I4521N, FWO, long stay abroad travel grant - grant number V438923N). D.R.-R. is supported by the Ministry of Universities through the “Pla de Recuperacio, Transformació i Resiliencia” and by the EU (NextGenerationEU), together with the Universitat de les Illes Balears. Furthermore, we thank Bartosz Prokop, Felix Nolet, and Arno Vanderbeke for valuable discussions and technical advice during the project.

## Author contributions statement

L.G. and L.P. conceived the study; L.P. conducted the experiments, aided by A.V.E., G.C.-M., L.G., and R.H.; N.F. developed the software; L.P. and N.F. analyzed the data; D.R.-R. and L.G. developed and analyzed the model; L.P., N.F., D.R.-R and L.G. prepared the figures and wrote the manuscript, with L.G. incorporating feedback from all authors.

## Declaration of generative AI and AI-assisted technologies in the writing process

During the preparation of this work the author(s) used ChatGPT in order to get feedback on the language of the written manuscript. We used the prompt “Check for grammatical errors in this text and indicate suggested corrections in bold.” After using this tool, the author(s) reviewed and edited the content as needed and take(s) full responsibility for the content of the publication.

## Movies

**Mov. 1 The nuclear compartment slows down the cycle duration in droplets of cycling extract**. Video of microtubule (MT) self-organization visualized by fluorescently labeled tubulin in cycling egg extract with and without demembranated sperm chromatin (SC). Corresponds to Figure 1.

**Mov. 2 Cycle duration positively correlates with nuclear-cytoplasmic ratio independent of initial nuclear size**. Cycling extracts were supplemented with different amounts of demembranated sperm chromatin (SC): 62.5, 250, and 500 SC*/µ*L. The videos show the nuclei (GFP-NLS) formed in the droplets of cell-free extracts. Corresponds to Figure 2.

**Mov. 3 Correlation between the cell cycle duration and the N/C ratio is preserved in *Xenopus laevis* cycling extract with added sperm chromatin**. Cycling extract were prepared using *X. laevis* eggs, and supplemented with sperm chromatin (SC) from the three frog species: (*X. laevis, X. tropicalis* or *X. borealis*). Corresponds to Figure 3.

**Mov. 4 Correlation between the cell cycle duration and the N/C ratio is preserved in *Xenopus laevis* cycling extract with added embryo chromatin**. Cycling extract were prepared using *X. laevis* eggs, and supplemented with stage 8 embryo chromatin (EC) from the three frog species: (*X. laevis, X. tropicalis* or *X. borealis*). Corresponds to Figure 3.

**Mov. 5 Correlation between the cell cycle duration and the N/C ratio is preserved in *Xenopus borealis* cycling extract with added sperm chromatin**. Cycling extract were prepared using *X. borealis* eggs, and supplemented with sperm chromatin (SC) from the three frog species: (*X. laevis, X. tropicalis* or *X. borealis*). Corresponds to Figure 3.

**Mov. 6 Wee1 inhibition disrupts the correlation between the cell cycle duration and the N/C ratio**. Cycling extracts were prepared using *X. laevis* eggs, and supplemented with *X. laevis* sperm chromatin (SC). Some droplets with extract were also supplemented with the inhibitor PD166285 targeting protein tyrosine kinase Wee1, which disrupted the correlation between the cell cycle duration and the N/C ratio. Corresponds to Figure 5.

**Mov. 7 Nuclear import inhibition disrupts the correlation between the cell cycle duration and the N/C ratio**. Cycling extracts were prepared using *X. laevis* eggs, and supplemented with *X. laevis* sperm chromatin (SC). Some droplets with extract were also supplemented with the inhibitor importazole targeting the nuclear import receptor importin-*—*, which disrupted the correlation between the cell cycle duration and the N/C ratio. Corresponds to Figure 5.

**Mov. 8 Inhibiting DNA replication does not affect the correlation between the cell cycle duration and the N/C ratio**. Cycling extracts were prepared using *X. laevis* eggs, and supplemented with *X. laevis* sperm chromatin (SC). Some droplets with extract were also supplemented with the inhibitor aphidicolin targeting DNA polymerase, which did not change the correlation between the cell cycle duration and the N/C ratio. Corresponds to Figure 6.

**Mov. 9 Inhibiting RNA polymerase II/III does not affect the correlation between the cell cycle duration and the N/C ratio**. Cycling extracts were prepared using *X. laevis* eggs, and supplemented with *X. laevis* sperm chromatin (SC). Some droplets with extract were also supplemented with the inhibitor *–*-amanitin targeting RNA polymerase II/III, which did not change the correlation between the cell cycle duration and the N/C ratio. Corresponds to Figure 6.

**Mov. 10 Inhibiting protein kinase Chk1 does not affect the correlation between the cell cycle duration and the N/C ratio**. Cycling extracts were prepared using *X. laevis* eggs, and supplemented with *X. laevis* sperm chromatin (SC). Some droplets with extract were also supplemented with the inhibitor SAR-020106 targeting protein kinase Chk1, which did not change the correlation between the cell cycle duration and the N/C ratio. Corresponds to Figure 6.

## References

1. Hirofumi Harashima, Nico Dissmeyer, and Arp Schnittger. Cell cycle control across the eukaryotic kingdom. Trends in cell biology, 23(7):345–356,.

2. David O Morgan. Cyclin-dependent kinases: engines, clocks, and microprocessors. Annual review of cell and developmental biology, 13(1):261–291,.

3. Jeffrey A Ubersax, Erika L Woodbury, Phuong N Quang, Maria Paraz, Justin D Blethrow, Kavita Shah, Kevan M Shokat, and David O Morgan. Targets of the cyclin-dependent kinase cdk1. Nature, 425(6960):859–864,.

4. Noah Dephoure, Chunshui Zhou, Judit Villén, Sean A Beausoleil, Corey E Bakalarski, Stephen J Elledge, and Steven P Gygi. A quantitative atlas of mitotic phosphorylation. Proceedings of the National Academy of Sciences, 105(31):10762–10767,.

5. Satoru Mochida, Sarah L Maslen, Mark Skehel, and Tim Hunt. Greatwall phosphorylates an inhibitor of protein phosphatase 2A that is essential for mitosis. Science (New York, N.Y.), 330(6011): 1670–1673, ec 2010.

6. Satoru Mochida and Tim Hunt. Protein phosphatases and their regulation in the control of mitosis. EMBO reports, 13(3):197–203, mar 2012.

7. Satoru Mochida, Scott Rata, Hirotsugu Hino, Takeharu Nagai, and Béla Novák. Two Bistable Switches Govern M Phase Entry. Current Biology, 26(24):3361–3367, ec 2016.

8. Julia Kamenz, Lendert Gelens, and James E. Ferrell. Bistable, biphasic regulation of pp2a-b55 accounts for the dynamics of mitotic substrate phosphorylation. Current Biology, 31(4):794–808.e6, 2021.

9. L Gelens, J Qian, M Bollen, and A T Saurin. The importance of kinase-phosphatase integration: Lessons from mitosis. Trends Cell Biol, 28:6–21,.

10. Anna Philpott and P Renee Yew. The xenopus cell cycle: an overview. Molecular biotechnology, 39:9–19,.

11. Andrew W Murray and Marc W Kirschner. Cyclin synthesis drives the early embryonic cell cycle. Nature, 339(6222):275–280, 1989.

12. A W Murray, M J Solomon, and M W Kirschner. The role of cyclin synthesis and degradation in the control of maturation promoting factor activity. Nature, 339(6222):280–286, may 1989.

13. Andrew W. Murray. Cell Cycle Extracts. Methods in Cell Biology, 36:581–605, 1991.

14. Jitender Bisht, Paige LeValley, Benjamin Noren, Ralph McBride, Prathamesh Kharkar, April Kloxin, Jesse Gatlin, and John Oakey. Light-inducible activation of cell cycle progression in xenopus egg extracts under microfluidic confinement. Lab on a Chip, 19(20):3499–3511,.

15. Y Masui and C L Markert. Cytoplasmic control of nuclear behavior during meiotic maturation of frog oocytes. J Exp Zool, 177:129–45,.

16. K Hara, P Tydeman, and M Kirschner. A cytoplasmic clock with the same period as the division cycle in Xenopus eggs. Proceedings of the National Academy of Sciences of the United States of America, 77(1):462–466, jan 1980.

17. J Gerhart, M Wu, and M Kirschner. Cell cycle dynamics of an m-phase-specific cytoplasmic factor in xenopus laevis oocytes and eggs. J Cell Biol, 98:1247–55,.

18. Frederick R. Cross and Eric D. Siggia. Shake it, don’t break it: Positive feedback and the evolution of oscillator design. Developmental Cell, 9(3):309–310, 2005.

19. James E Jr Ferrell and Sang Hoon Ha. Ultrasensitivity part III: cascades, bistable switches, and oscillators. Trends in biochemical sciences, 39(12):612–618, ec 2014.

20. James E Jr Ferrell and Sang Hoon Ha. Ultrasensitivity part II: multisite phosphorylation, stoichiometric inhibitors, and positive feedback. Trends in biochemical sciences, 39(11):556–569, nov 2014.

21. Jolan De Boeck, Jan Rombouts, and Lendert Gelens. A modular approach for modeling the cell cycle based on functional response curves. PLOS Computational Biology, 17(8):1–39,.

22. Tony Yu-Chen Tsai, Yoon Sup Choi, Wenzhe Ma, Joseph R Pomerening, Chao Tang, and James E Jr Ferrell. Robust, tunable biological oscillations from interlinked positive and negative feedback loops. Science (New York, N.Y.), 321(5885):126–129, jul 2008.

23. Pedro Parra-Rivas, Daniel Ruiz-Reynés, and Lendert Gelens. Cell cycle oscillations driven by two interlinked bistable switches. Molecular Biology of the Cell, 34(6):ar56,.

24. Xianrui Cheng and James E Ferrell Jr. Spontaneous emergence of cell-like organization in xenopus egg extracts. Science, 366(6465):631–637,.

25. Timothy J Mitchison and Christine M Field. Self-organization of cellular units. Annual review of cell and developmental biology, 37:23–41,.

26. Pierre-Yves Gires, Mithun Thampi, Sebastian W Krauss, and Matthias Weiss. Exploring generic principles of compartmentalization in a developmental in vitro model. Development, 150(3): dev200851,.

27. Jan Rombouts and Lendert Gelens. Dynamic bistable switches enhance robustness and accuracy of cell cycle transitions. PLoS computational biology, 17(1):e1008231,.

28. Gembu Maryu and Qiong Yang. Nuclear-cytoplasmic compartmentalization of cyclin b1-cdk1 promotes robust timing of mitotic events. Cell Reports, 41(13):111870, 2022.

29. Felix E Nolet, Alexandra Vandervelde, Arno Vanderbeke, Liliana Piñeros, Jeremy B Chang, and Lendert Gelens. Nuclei determine the spatial origin of mitotic waves. eLife, 9:e52868, may 2020.

30. Oshri Afanzar, Garrison K Buss, Tim Stearns, and Jr Ferrell, James E. The nucleus serves as the pacemaker for the cell cycle. eLife, 9:e59989, ec 2020.

31. O. Puls*, D. Ruiz-Reynés*, F. Tavella, M. Jin, Y. Kim, L. Gelens*, and Q. Yang*. Spatial heterogeneity accelerates phase-to-trigger wave transitions in frog egg extracts. Nature Communications, 15 (10455),.

32. Silvia DM Santos, Roy Wollman, Tobias Meyer, and James E Ferrell. Spatial positive feedback at the onset of mitosis. Cell, 149(7):1500–1513,.

33. Ye Guan, Zhengda Li, Shiyuan Wang, Patrick M Barnes, Xuwen Liu, Haotian Xu, Minjun Jin, Allen P Liu, and Qiong Yang. A robust and tunable mitotic oscillator in artificial cells. eLife, 7:e33549, apr 2018.

34. Jeremy B Chang and James E Ferrell. Robustly cycling xenopus laevis cell-free extracts in teflon chambers. Cold Spring Harbor Protocols, 2018(8):pdb–prot097212,.

35. Angela Flavia Serpico, Francesco Febbraro, Caterina Pisauro, and Domenico Grieco. Compartmentalized control of cdk1 drives mitotic spindle assembly. Cell Reports, 38(4):110305, 2022.

36. Graham A. Anderson, Lendert Gelens, Julie C. Baker, and James E. Ferrell. Desynchronizing embryonic cell division waves reveals the robustness of xenopus laevis development. Cell Reports, 21 (1):37–46, 2017.

37. John Newport and Marc Kirschner. A major developmental transition in early xenopus embryos: I. characterization and timing of cellular changes at the midblastula stage. Cell, 30(3):675–686,.

38. Noriyuki Satoh. ‘Metachronous’ cleavage and initiation of gastrulation in amphibian embryos. Development, Growth Differentiation, 19(2):111–117, jan 1977.

39. Rebecca Heald, Michael McLoughlin, and Frank McKeon. Human wee1 maintains mitotic timing by protecting the nucleus from cytoplasmically activated cdc2 kinase. Cell, 74(3):463–474,.

40. Véronique Baldin and Bernard Ducommun. Subcellular localisation of human wee1 kinase is regulated during the cell cycle. Journal of Cell Science, 108(6):2425–2432, 06 1995.

41. Jonathan D. Moore, Jing Yang, Ray Truant, and Sally Kornbluth. Nuclear Import of Cdk/Cyclin Complexes: Identification of Distinct Mechanisms for Import of Cdk2/Cyclin E and Cdc2/Cyclin B1. Journal of Cell Biology, 144(2):213–224, 01 1999.

42. Hanna Salman, Asmahan Abu-Arish, Shachar Oliel, Avraham Loyter, Joseph Klafter, Rony Granek, and Michael Elbaum. Nuclear localization signal peptides induce molecular delivery along microtubules. Biophysical Journal, 89(3):2134–2145, 2005.

43. Olivier Gavet and Jonathon Pines. Activation of cyclin B1–Cdk1 synchronizes events in the nucleus and the cytoplasm at mitosis. Journal of Cell Biology, 189(2):247–259, 04 2010.

44. Sahla Syed, Henry Wilky, João Raimundo, Bomyi Lim, and Amanda A Amodeo. The nuclear to cytoplasmic ratio directly regulates zygotic transcription in <i>Drosophila</i> through multiple modalities. Proceedings of the National Academy of Sciences, 118(14):e2010210118, apr 2021.

45. Predrag Jevtić and Daniel L Levy. Both nuclear size and dna amount contribute to midblastula transition timing in xenopus laevis. Scientific Reports, 7:7908, 12 2017.

46. Adam M Session, Yoshinobu Uno, Taejoon Kwon, Jarrod A Chapman, Atsushi Toyoda, Shuji Takahashi, Akimasa Fukui, Akira Hikosaka, Atsushi Suzuki, Mariko Kondo, et al. Genome evolution in the allotetraploid frog xenopus laevis. Nature, 538(7625):336–343,.

47. Maiko Kitaoka, Rebecca Heald, and Romain Gibeaux. Spindle assembly in egg extracts of the marsabit clawed frog, xenopus borealis. Cytoskeleton, 75(6):244–257,.

48. Romain Gibeaux, Kelly Miller, Rachael Acker, Taejoon Kwon, and Rebecca Heald. Xenopus hybrids provide insight into cell and organism size control. Frontiers in physiology, 9:1758,.

49. Romain Gibeaux and Rebecca Heald. Generation of Xenopus Haploid, Triploid, and Hybrid Embryos, pages 303–315. Springer New York, New York, NY, 2019.

50. Kai Yuan, Charles A. Seller, Antony W. Shermoen, and Patrick H. O’Farrell. Timing the drosophila mid-blastula transition: A cell cycle-centered view. Trends in Genetics, 32(8):496–507, 2016.

51. Qiong Yang and James E. Ferrell. The Cdk1-APC/C cell cycle oscillator circuit functions as a time-delayed, ultrasensitive switch. Nature Cell Biology, 15(5):519–525, may 2013.

52. Joseph R. Pomerening, Eduardo D. Sontag, and James E. Ferrell. Building a cell cycle oscillator: Hysteresis and bistability in the activation of Cdc2. Nature Cell Biology, 5(4):346–351, apr 2003.

53. Wei Sha, Jonathan Moore, Katherine Chen, Antonio D Lassaletta, Chung-Seon Yi, John J Tyson, and Jill C Sible. Hysteresis drives cell-cycle transitions in Xenopus laevis egg extracts. Proceedings of the National Academy of Sciences, 100(3):975–980, 2003.

54. Scott Rata, Maria F Suarez Peredo Rodriguez, Stephy Joseph, Nisha Peter, Fabio Echegaray Iturra, Fengwei Yang, Anotida Madzvamuse, Jan G Ruppert, Kumiko Samejima, Melpomeni Platani, Monica Alvarez-Fernandez, Marcos Malumbres, William C Earnshaw, Bela Novak, and Helfrid Hochegger. Two Interlinked Bistable Switches Govern Mitotic Control in Mammalian Cells. Current Biology, 28(23):3824–3832.e6, 2018.

55. Yuli Wang, Jun Li, Robert N Booher, Alan Kraker, Theodore Lawrence, Wilbur R Leopold, and Yi Sun. Radiosensitization of p53 mutant cells by pd0166285, a novel g2 checkpoint abrogator. Cancer research, 61(22):8211–8217,.

56. Osamu Hashimoto, Masako Shinkawa, Takuji Torimura, Toru Nakamura, Karuppaiyah Selvendiran, Masaharu Sakamoto, Hironori Koga, Takato Ueno, and Michio Sata. Cell cycle regulation by the wee1 inhibitor pd0166285, pyrido [2,3-d] pyimidine, in the b16 mouse melanoma cell line. BMC Cancer, 6:292, 2006.

57. Amnon Harel, Rene C Chan, Aurelie Lachish-Zalait, Ella Zimmerman, Michael Elbaum, and Douglass J Forbes. Importin — negatively regulates nuclear membrane fusion and nuclear pore complex assembly. Molecular biology of the cell, 14(11):4387–4396,.

58. Amnon Harel and Douglass J Forbes. Importin beta: conducting a much larger cellular symphony. Molecular cell, 16(3):319–330,.

59. John Newport and Marc Kirschner. A major developmental transition in early xenopus embryos: I. characterization and timing of cellular changes at the midblastula stage. Cell, 30(3):675–686, 1982.

60. John Newport and Marc Kirschner. A major developmental transition in early xenopus embryos: II. control of the onset of transcription. Cell, 30(3):687–696, 1982.

61. Bruce A Edgar, Caroline P Kiehle, and Gerold Schubiger. Cell cycle control by the nucleo-cytoplasmic ratio in early Drosophila development. Cell, 44(2):365–372, 1986.

62. Hui Chen, Lily C Einstein, Shawn C Little, and Matthew C Good. Spatiotemporal Patterning of Zygotic Genome Activation in a Model Vertebrate Embryo. Developmental Cell, 49(6):852–866.e7, 2019.

63. Hiroko Heijo, Sora Shimogama, Shuichi Nakano, Anna Miyata, Yasuhiro Iwao, and Yuki Hara. Dna content contributes to nuclear size control in xenopus laevis. Molecular Biology of the Cell, 31 (24):2703–2717,.

64. Jeffrey A Farrell and Patrick H O’Farrell. From egg to gastrula: how the cell cycle is remodeled during the Drosophila mid-blastula transition. Annual review of genetics, 48:269–294, 2014.

65. Maomao Zhang, Jennifer Skirkanich, Michael A. Lampson, and Peter S. Klein. Cell cycle remodeling and zygotic gene activation at the midblastula transition. 953:441–487, 2017.

66. Alexey G. Desnitskiy. Cell cycles during early steps of amphibian embryogenesis: A review. 173:100–103, nov 2018.

67. Amanda A Amodeo, David Jukam, Aaron F Straight, and Jan M Skotheim. Histone titration against the genome sets the dna-to-cytoplasm threshold for the xenopus midblastula transition. Proceedings of the National Academy of Sciences, 112(10):E1086–E1095,.

68. Shruthi Balachandra, Sharanya Sarkar, and Amanda A. Amodeo. The nuclear-to-cytoplasmic ratio: Coupling dna content to cell size, cell cycle, and biosynthetic capacity. Annual Review of Genetics, 56(1):165–185,. PMID: 35977407.

69. Yuki Shindo and Amanda A Amodeo. Excess histone H3 is a competitive Chk1 inhibitor that controls cell-cycle remodeling in the early Drosophila embryo. Current Biology, 31(12):2633–2642.e6, 2021.

70. Yuki Shindo and Amanda A Amodeo. Modeling the role for nuclear import dynamics in the early embryonic cell cycle. Biophysical Journal, 120(19):4277–4286,.

71. Elze C Boterenbrood, Jennifer M Narraway, and Koki Hara. Duration of cleavage cycles and asymmetry in the direction of cleavage waves prior to gastrulation inXenopus laevis. Wilhelm Roux’s archives of developmental biology, 192(5):216–221, 1983.

72. Nicolas Olivier, Miguel A Luengo-Oroz, Louise Duloquin, Emmanuel Faure, Thierry Savy, Israël Veilleux, Xavier Solinas, Delphine Débarre, Paul Bourgine, Andrés Santos, Nadine Peyriéras, and Emmanuel Beaurepaire. Cell lineage reconstruction of early zebrafish embryos using label-free nonlinear microscopy. Science (New York, N.Y.), 329(5994):967–971, aug 2010.

73. Shruthi Balachandra, Sharanya Sarkar, and Amanda A. Amodeo. The nuclear-to-cytoplasmic ratio: Coupling dna content to cell size, cell cycle, and biosynthetic capacity. Annual Review of Genetics, 56:165–185, 11 2022.

74. Gabriel E. Neurohr, Rachel L. Terry, Jette Lengefeld, Megan Bonney, Gregory P. Brittingham, Fabien Moretto, Teemu P. Miettinen, Laura Pontano Vaites, Luis M. Soares, Joao A. Paulo, J. Wade Harper, Stephen Buratowski, Scott Manalis, Folkert J. van Werven, Liam J. Holt, and Angelika Amon. Excessive cell growth causes cytoplasm dilution and contributes to senescence. Cell, 176: 1083–1097.e18, 2 2019.

75. Michael C. Lanz, Evgeny Zatulovskiy, Matthew P. Swaffer, Lichao Zhang, Ilayda Ilerten, Shuyuan Zhang, Dong Shin You, Georgi Marinov, Patrick McAlpine, Joshua E. Elias, and Jan M. Skotheim. Increasing cell size remodels the proteome and promotes senescence. Molecular Cell, 82:3255–3269.e8, 9 2022.

76. Shicong Xie, Matthew Swaffer, and Jan M. Skotheim. Eukaryotic cell size control and its relation to biosynthesis and senescence. Annual Review of Cell and Developmental Biology, 38:291–319, 10 2022.

77. Ling Cheng, Jingyuan Chen, Yidi Kong, Ceryl Tan, Ran Kafri, and Mikael Björklund. Size-scaling promotes senescence-like changes in proteome and organelle content. bioRxiv, page 2021.08.05.455193,.

78. Sandhya Manohar and Gabriel E. Neurohr. Too big not to fail: emerging evidence for size-induced senescence. The FEBS Journal, 291:2291–2305, 6 2024.

79. Aresu Sadeghi Shoreh Deli, Sonja Scharf, Yvonne Steiner, Julia Bein, Martin Leo Hansmann, and Sylvia Hartmann. 3d analyses reveal t cells with activated nuclear features in t-cell/histiocyte-rich large b-cell lymphoma. Modern Pathology, 35:1431–1438, 10 2022.

80. Niki Karachaliou, Sara Pilotto, Chiara Lazzari, Emilio Bria, Filippo de Marinis, and Rafael Rosell. Cellular and molecular biology of small cell lung cancer: an overview. Translational Lung Cancer Research, 5:2–15, 2016.

81. Jose I. De las Heras, Dzmitry G. Batrakou, and Eric C. Schirmer. Cancer biology and the nuclear envelope: A convoluted relationship. Seminars in Cancer Biology, 23:125–137, 4 2013.

82. Hsu-Cheng Huang, Shu-Jen Chiang, Shu-Han Wen, Pei-Jung Lee, Huei-Wen Chen, Yang-Fang Chen, and Chen-Yuan Dong. Three-dimensional nucleus-to-cytoplasm ratios provide better discrimination of normal and lung adenocarcinoma cells than in two dimensions. 10.1117/1.JBO.24.8.080502, 24:080502, 8 2019.

83. Abin Biswas, Omar Muñoz, Kyoohyun Kim, Carsten Hoege, Benjamin M Lorton, David Shechter, Jochen Guck, Vasily Zaburdaev, and Simone Reber. Conserved nucleocytoplasmic density homeostasis drives cellular organization across eukaryotes. bioRxiv, pages 2023–09,.

84. Durrieu Lucía, Bush Alan, Grande Alicia, Johansson Rikard, Janzén David, Andrea Katz, Gunnar Cedersund, and Colman-Lerner Alejandro. Characterization of cell-to-cell variation in nuclear transport rates and identification of its sources. Iscience, 26(1),.

85. Andrew W. Murray. Chapter 30 cell cycle extracts. In Brian K. Kay and H. Benjamin Peng, editors, Xenopus laevis: Practical Uses in Cell and Molecular Biology, volume 36 of Methods in Cell Biology, pages 581–605. Academic Press,.

86. Coral Y Zhou, Bastiaan Dekker, Ziyuan Liu, Hilda Cabrera, Joel Ryan, Job Dekker, and Rebecca Heald. Mitotic chromosomes scale to nuclear-cytoplasmic ratio and cell size in Xenopus. eLife, 12: e84360, apr 2023.

87. Christine M. Field, Martin Wühr, Graham A. Anderson, Hao Yuan Kueh, Devin Strickland, and Timothy J. Mitchison. Actin behavior in bulk cytoplasm is cell cycle regulated in early vertebrate embryos. Journal of Cell Science, 124(12):2086–2095, 06 2011.

88. Marius Pachitariu and Carsen Stringer. Cellpose 2.0: how to train your own model. Nature Methods, 19(12):1634–1641,.

89. Guillaume Witz and Peter Sobolewski. Napari-serialcellpose, 2022.

90. Nicholas Sofroniew, Talley Lambert, Kira Evans, Juan Nunez-Iglesias, Grzegorz Bokota, Philip Winston, Gonzalo Peña-Castellanos, Kevin Yamauchi, Matthias Bussonnier, Draga Doncila Pop, Ahmet Can Solak, Ziyang Liu, Pam Wadhwa, Alister Burt, Genevieve Buckley, Andrew Sweet, Lukasz Migas, Volker Hilsenstein, Lorenzo Gaifas, Jordão Bragantini, Jaime Rodríguez-Guerra, Hector Muñoz, Jeremy Freeman, Peter Boone, Alan Lowe, Christoph Gohlke, Loic Royer, Andrea Pierré, Hagai Har-Gil, and Abigail McGovern. Napari: multi-dimensional image viewer for python, 2022.

91. Ron Milo and Rob Phillips. Cell biology by the numbers. Garland Science,.

92. Mohit Kumar, Mario S Mommer, and Victor Sourjik. Mobility of cytoplasmic, membrane, and dna-binding proteins in escherichia coli. Biophysical journal, 98(4):552–559,.

93. Predrag Jevtić and Daniel L Levy. Nuclear size scaling during xenopus early development contributes to midblastula transition timing. Current Biology, 25(1):45–52,.

94. Nikita Frolov, Liliana Piñeros, and Lendert Gelens. https://gitlab.kuleuven.be/gelenslab/publications/nuclear-compartment, 2024.

95. Daniel Ruiz-Reynès and Lendert Gelens. https://gitlab.kuleuven.be/gelenslab/publications/compartmentmodel, 2024.

96. Jeremy B Chang and James E Ferrell Jr. Mitotic trigger waves and the spatial coordination of the Xenopus cell cycle. Nature, 500(7464):603–607, 2013.

97. Jan Rombouts, Alexandra Vandervelde, and Lendert Gelens. Delay models for the early embryonic cell cycle oscillator. PLoS ONE, 13(3), mar 2018.

98. Matthias Eibauer, Mauro Pellanda, Yagmur Turgay, Anna Dubrovsky, Annik Wild, and Ohad Medalia. Structure and gating of the nuclear pore complex. Nature communications, 6(1):7532,.

