## Supplemental figures for "The nuclear-cytoplasmic ratio controls the cell cycle period in compartmentalized frog egg extract"

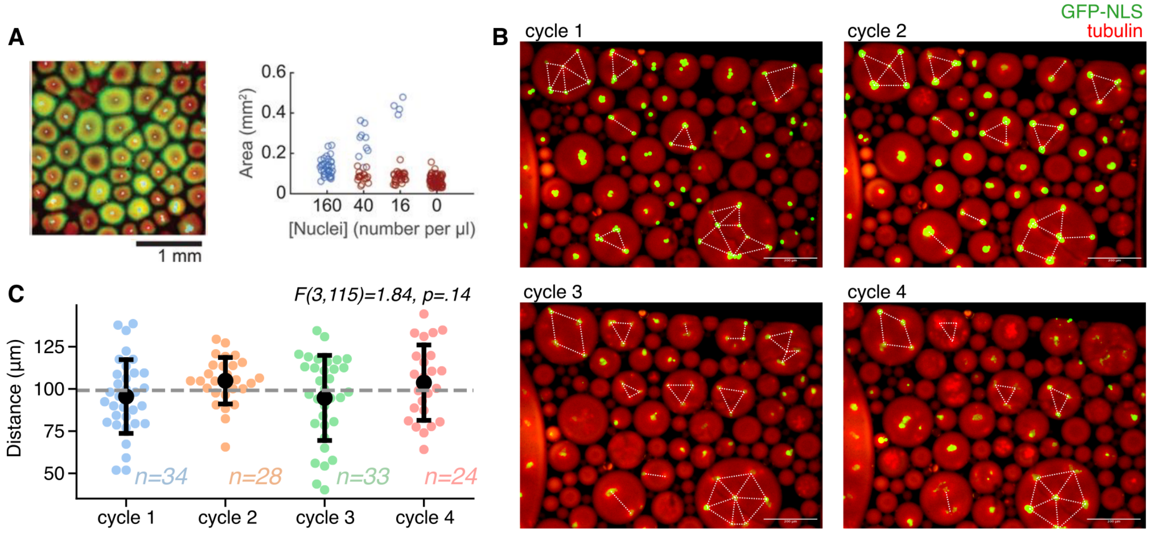


Fig. S1: **Compartmentalization of cell-free extract supplemented with SC in larger droplets. A.** (Left) A micrograph of compartmentalization in bulk cell-free extract supplemented with 160 nuclei/$\mu l$ arrested in the interphase. Tubulin, ER, and nuclei are stained. (Right) Distribution of compartment areas. The figure is taken from (24). **B.** Typical micrographs of droplets supplemented with cycling cell-free extracts at four successive cycles imaged in our study. Tubulin and nuclei are stained. Larger droplets (around 150 $\mu m$ and larger in diameter) contain multiple nuclei/compartments. White dashed lines display measured internuclear distances. Scale bars are 200 $\mu m$. **C.** Variation of inter-nuclei distance against cycle number (mean±SD). The dashed gray line indicates the general mean. One-way ANOVA was performed.

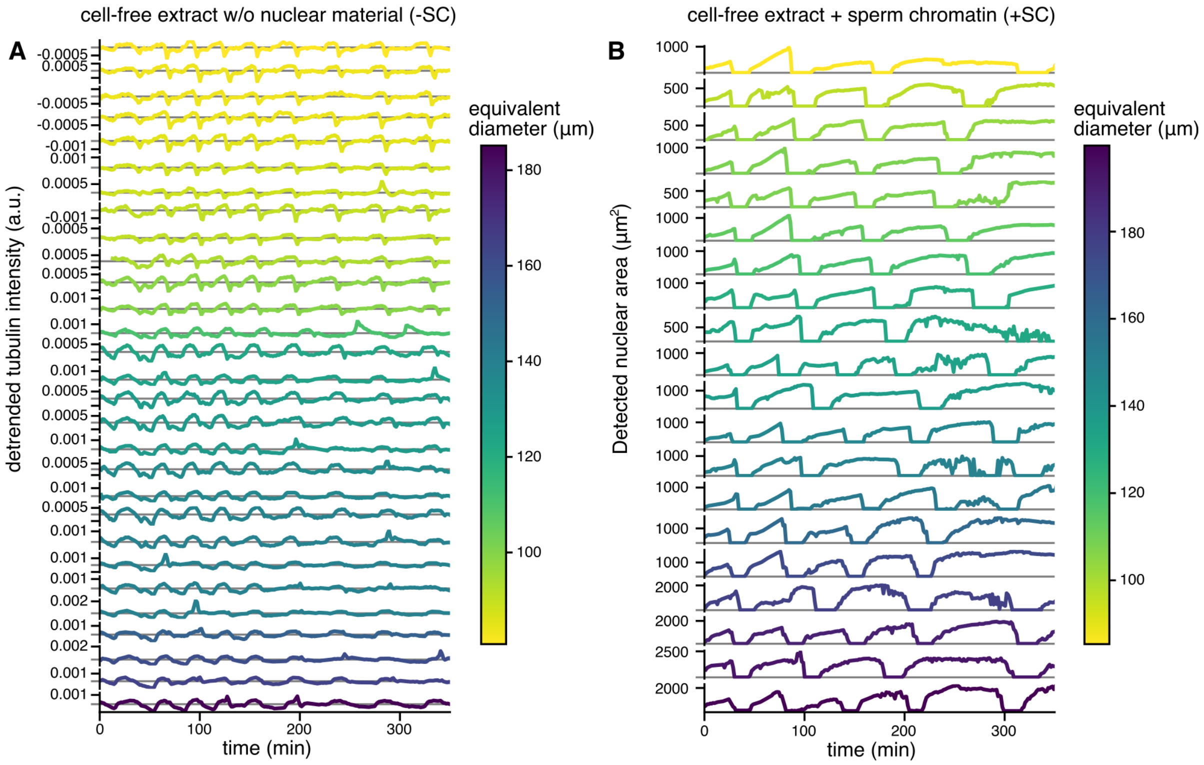


Fig. S2: **Time traces of cell cycle oscillations measured across multiple droplets of cell-free extract in the experiment from Fig. 1. A.** Time traces of the fluorescence intensity of labeled tubulin in a.u. measured in $n = 29$ droplets of cell-free extract without nuclear material (-SC). **B.** Time traces of the nuclear area in $\mu$m$^2$ measured by the Cellpose 2.0 algorithm in $n = 20$ droplets of cell-free extract supplemented with sperm chromatin (+SC). The curves in each panel are color-coded by the droplet size (equivalent diameter in $\mu$m).

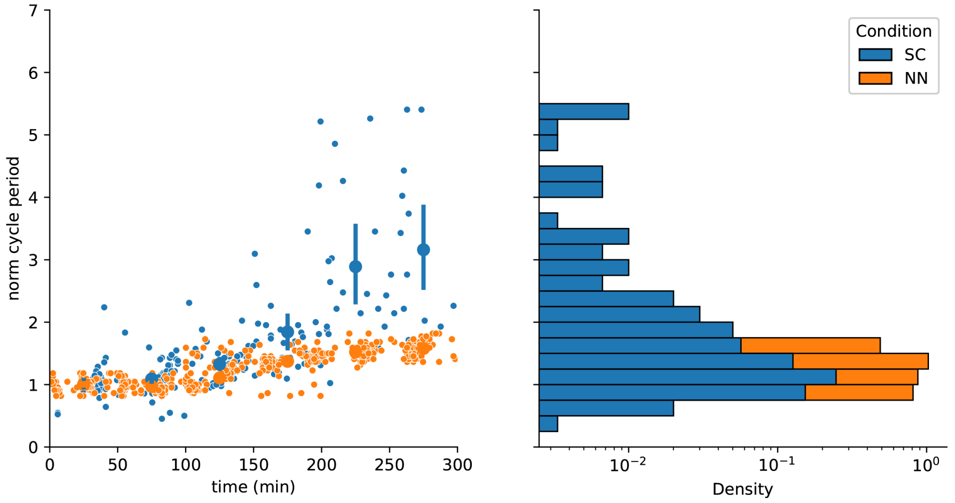


Fig. S3: **Slowing down of the cell cycle is more prominent in the presence of nuclei compared to the no-nuclei condition.** Semi-log plot of temporal variation of the cycle duration in cycling extract after adding demembranated SC. This is similar to Figure 1G, but here, the cycle duration is normalized to the initial cell cycle period, allowing us to compare the fold change over time. Mean±SD cycle duration in droplets with and without nuclei are shown by hollow circles with error bars. Histograms to the right show the distribution of the aggregate cycle duration for both conditions. Solid circles and error bars indicate the median and interquartile range, respectively.

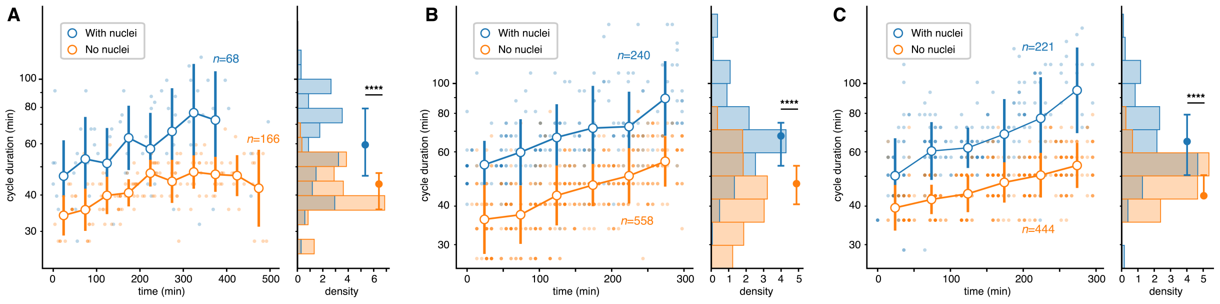


Fig. S4: **Slowing down of the cell cycle is more prominent in the presence of nuclei compared to the no-nuclei condition in 3 independent measurements.** In each subplot, the left panel presents a semi-log plot of temporal variation of the cycle duration for cycling extract with and without added sperm nuclei (hollow circles show mean±SD obtained by binning). Histograms in the right panels show the distributions of the aggregate cycle duration for both conditions. Solid circles and error bars indicate the median and interquartile range, respectively. '****' indicates the significance level of $p < .001$ via the Mann-Whitney test.

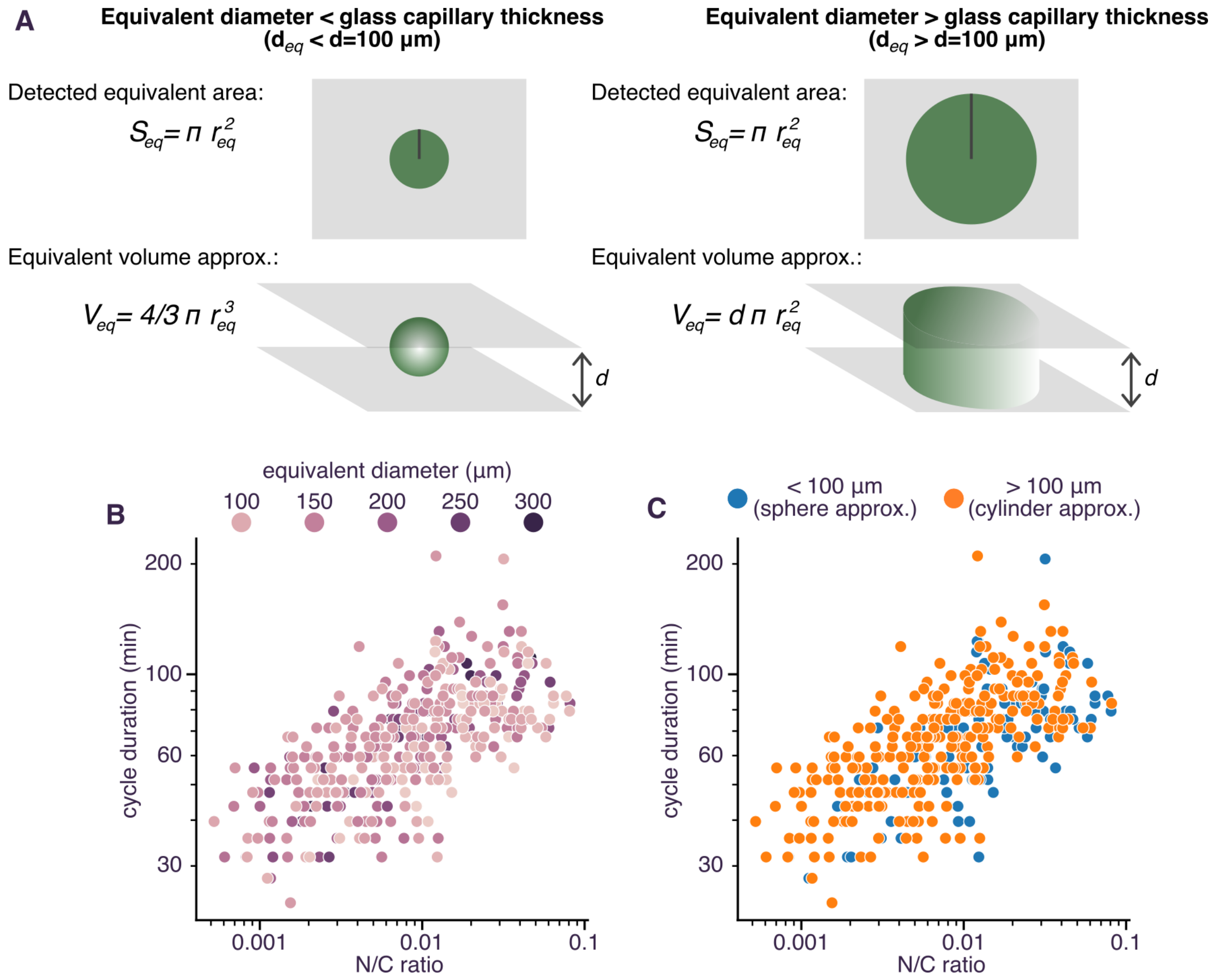


Fig. S5: **Rescaling detected droplet areas to volumes approximating droplets with either equivalent spheres or cylinders.** **A.** Schematic illustration of droplet geometry approximation for droplets of different sizes. **B, C.** Log-log plot of cell cycle duration versus N/C ratio equivalent to Fig. 2C. The scatterplot in panel B is color-coded with the droplet's equivalent diameter. Panel C shows the fractions of data obtained by spherical (blue) and cylindrical (orange) approximations of droplet geometry.

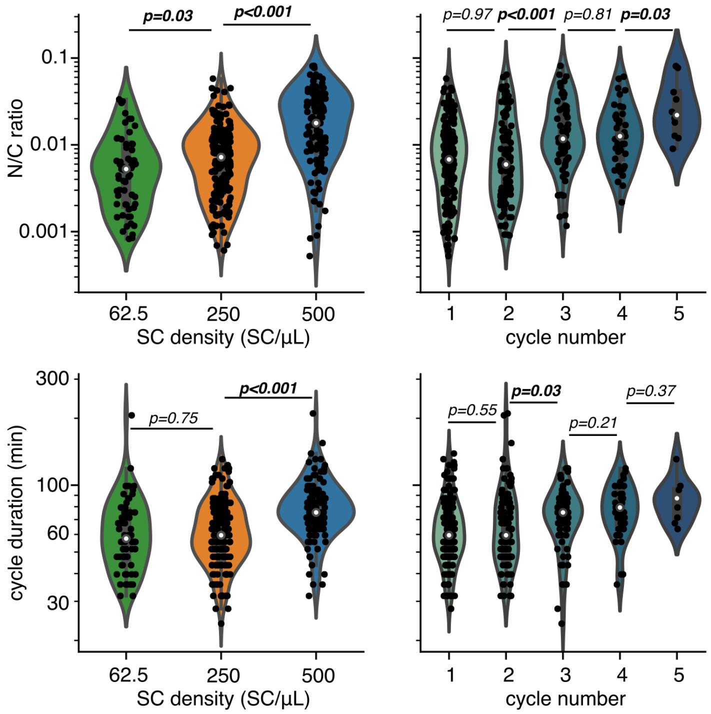


Fig. S6: Variation of the nuclear-cytoplasmic ratio (top) and cycle duration (bottom) versus the sperm chromatin density and the number of cycles. The Mann-Whitney test was performed for pair-wise comparisons.

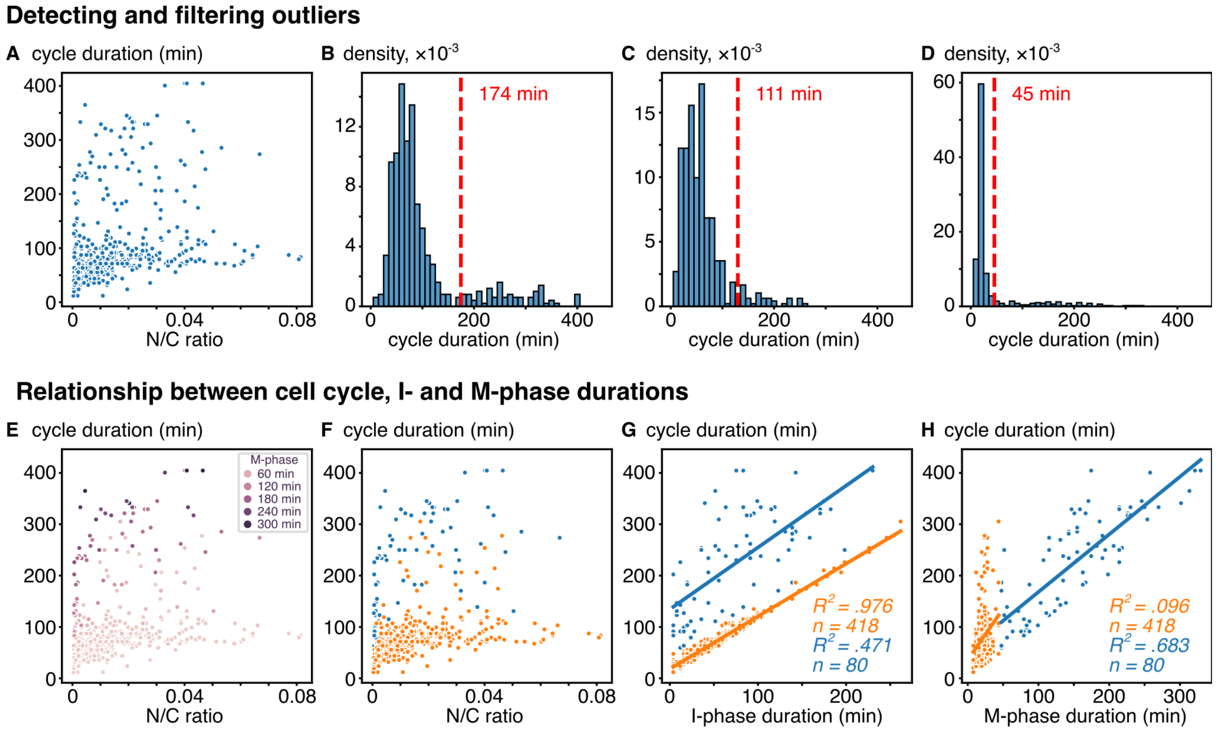


Fig. S7: **Outliers of the cycle duration distribution is associated with elongated M-phase. A.** Raw variance of the cycle duration versus N/C ratio. Density function of the cycle duration (**B**), I-phase duration (**C**), and M-phase duration (**D**) with a red dashed line indicating the outlier threshold. **E.** Raw variance of the cycle duration versus N/C ratio color-coded with the M-phase duration. **F.** Raw variance of the cycle duration versus N/C ratio with the M-phase outliers colored blue and normal population colored orange. Variance of cycle duration versus I-phase duration (**G**) and M-phase duration (**H**) for normal population and outliers. Lines of respective color indicate fitted linear models. Legends report the quality of the fit and sample size.

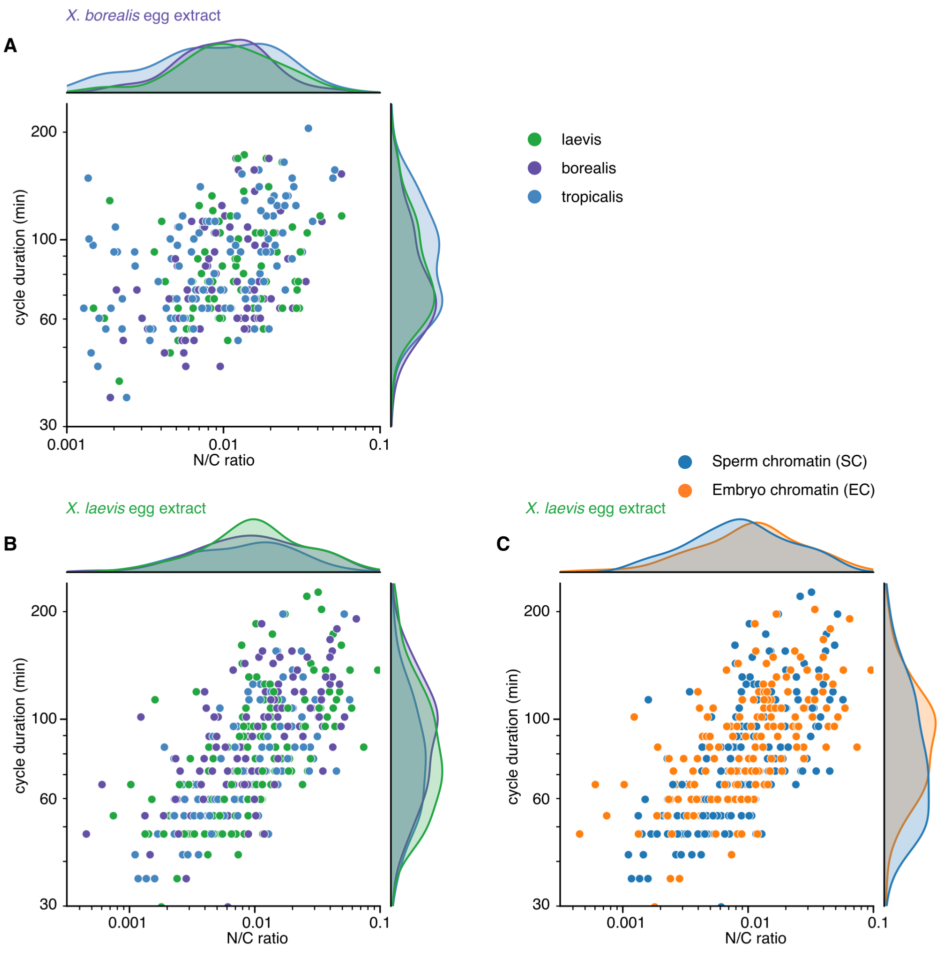


Fig. S8: **The cell cycle duration scales with the nuclear-to-cytoplasmic ratio in egg extract from various frog species. A.** Log-log plot of cell cycle duration versus N/C ratio with the histograms in egg extract from *X. borealis* supplemented with sperm chromatin and color-coded based on the frog species: *X. laevis* in green, *X. borealis* in purple, and *X. tropicalis* in blue. Log-log plot of cell cycle duration versus N/C ratio with the histograms in egg extract from *X. laevis* supplemented with sperm chromatin or stage 8 embryo nuclei from the same three frog species **B.** Color-coded based on the frog species and **C.** Color-coded based on the type of nuclear material: sperm chromatin in blue and embryo nuclei in orange.

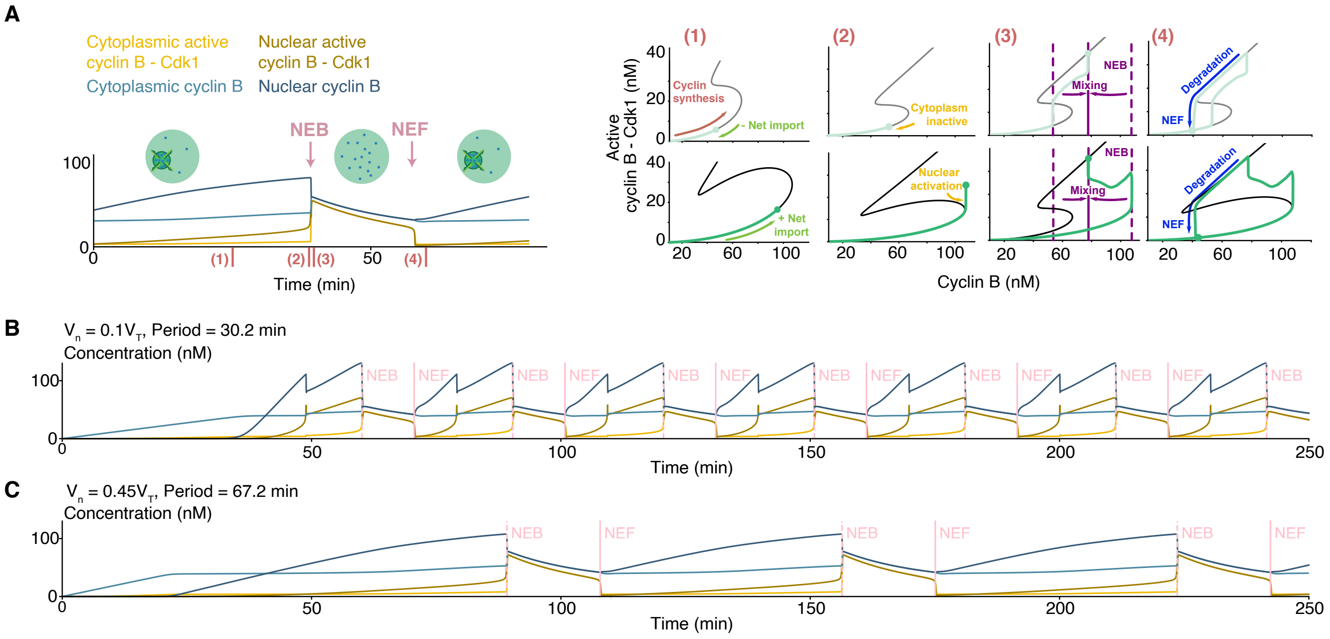


Fig. S9: **Mathematical modeling shows that nuclear compartmentalization can slow down the cell cycle period. A** Time series and phase plane representation of the dynamics of the cytoplasmic (light green) and nuclear (green) compartments in different phases of the cell cycle. According to NEB, S-shaped switches in the cytoplasm (gray) and nucleus (black) change dynamically. **B-C** Time series for two values of the nuclear volume as indicated by the labels. The concentration of cyclin b (blue) and active cyclin B - Cdk1 (yellow) in the nucleus (cytoplasm) is shown with continuous (dashed) lines. NEF (NEB) are indicated with vertical continuous (dashed) lines.

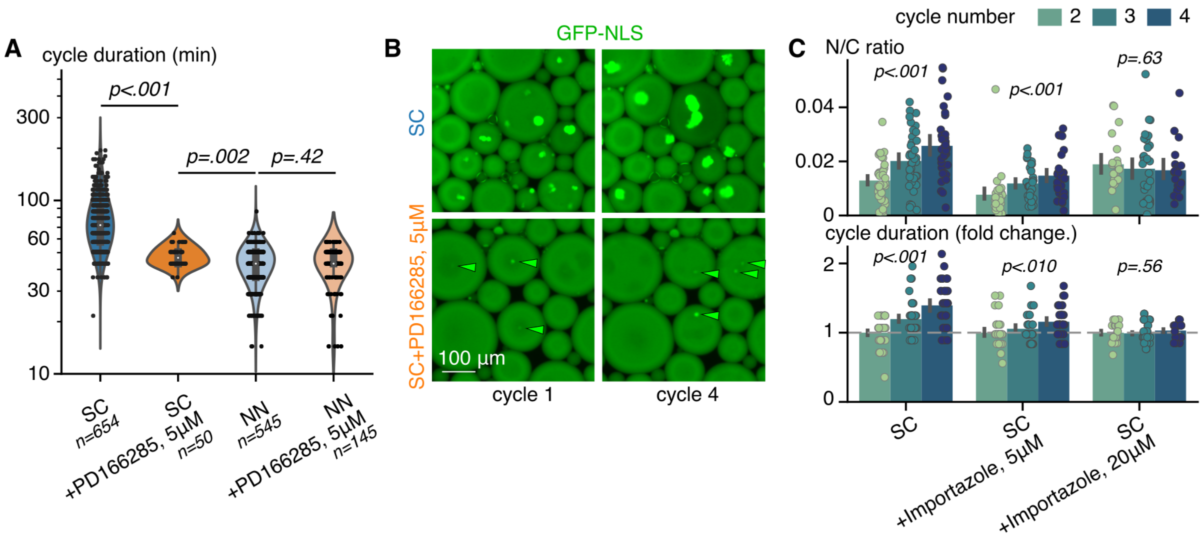


Fig. S10: **Supplemental data to Wee1 kinase and nuclear import inhibition experiments. A.** Comparison of cycle duration in extracts with (SC) and without (NN) added SC, and with or without Wee1 inhibition by PD166285. The Bonferroni-corrected Mann-Whitney U-test was performed for pair-wise comparisons. **B.** Micrographs of GFP-NLS show the nuclei formed at cycles 1 and 4 in the control extract (SC, top panels) and extract treated with Wee1 inhibitor (SC+PD166285, bottom panels). Arrowheads indicate nuclei in the PD166285-treated condition. Variation of N/C volume ratio (mean±SD, top) and fold increase in cycle duration (means±SD, bottom) during cycles 2-4 under different concentrations of importazole. The Kruskal-Wallis test was performed.

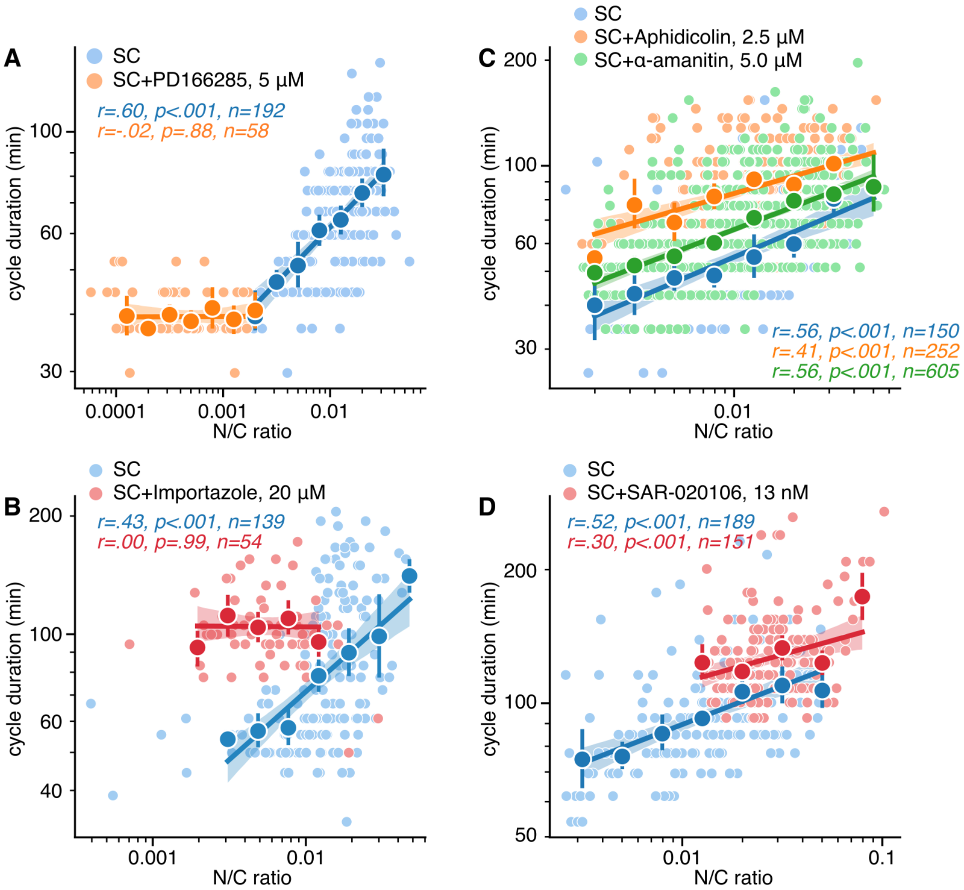


Fig. S11: **Independently repeated experiments treating cell-free extracts with various inhibitors.** Panels show the log-log plots of the cycle duration versus the N/C ratio under treatment with Wee1 inhibitor PD166285, 5 $\mu$M (**A**); with nuclear import inhibitor Importazole, 20 $\mu$M (**B**); with DNA replication inhibitor Aphidicolin, 2.5 $\mu$M, and DNA transcription inhibitor $\alpha$-amanitin, 5 $\mu$M (**C**); with Chk1 inhibitor SAR-020106, 13 nM (**D**). The pale circles display the data; the dark circles show the means obtained by binning the data, and the error bars indicate respective 95% confidence intervals. Solid lines represent the linear fit on the log-log plane. **B.** Typical time traces of labeled tubulin fluorescence intensity (red) and N/C ratio (green) in respective experimental conditions. **C.** The time moment of the first cell cycle detection versus experimental condition.

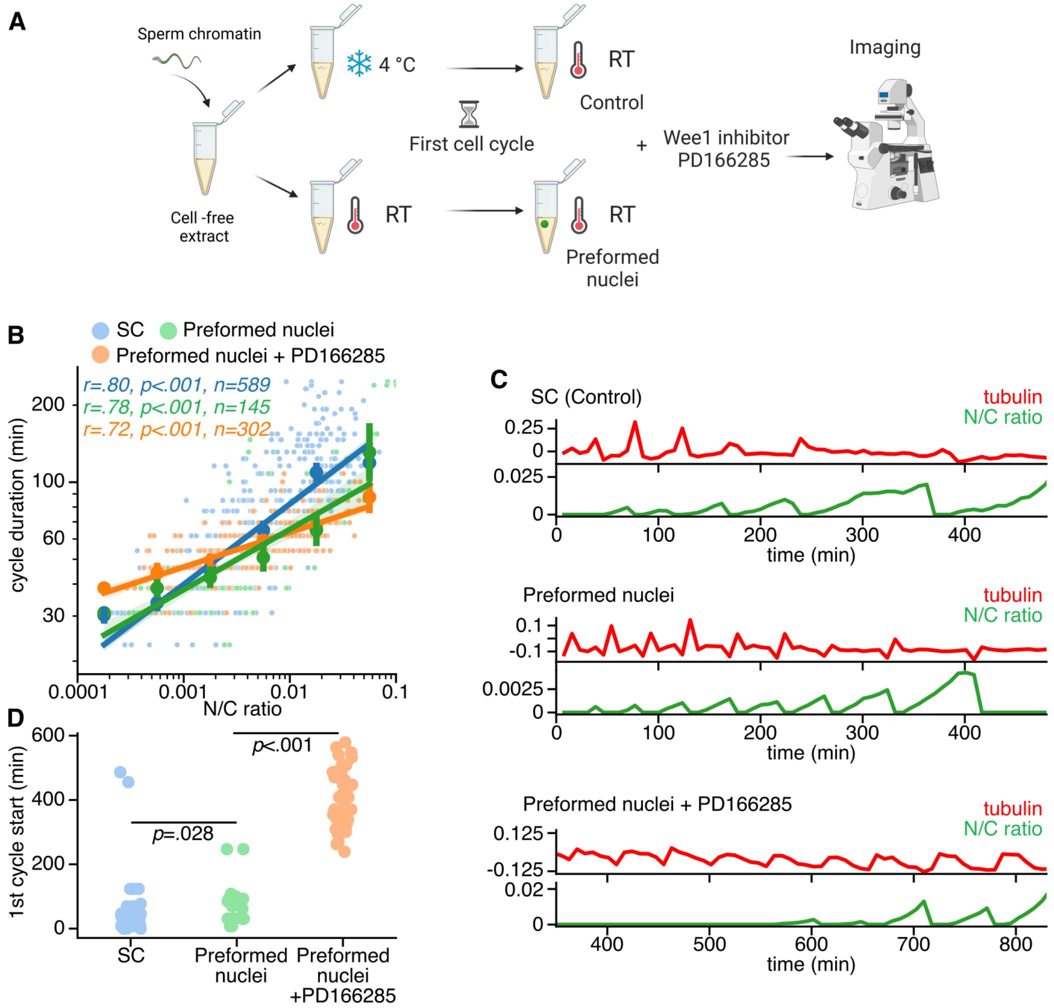


Fig. S12: **Supplemental data to Wee1 kinase inhibition experiment with preformed nuclei. A.** Scheme showing the experimental design testing the effect of Wee1 inhibition in the extracts with preformed nuclei. **B.** Log-log plot of cell cycle duration versus N/C ratio for control (SC, blue), Preformed nuclei (green), and Preformed nuclei+PD166285 (orange) conditions. The dark circles show the means obtained by binning the data, and the error bars indicate respective 95% confidence intervals. Solid lines represent the linear fit on the log-log plane. **C.** Typical time series of fluorescent tubulin intensity (red) and N/C volume ratio (green) measured in the cycling extract droplets in control (SC-supplemented), Wee1 inhibition (SC + PD166285), and Wee1 inhibited with preformed nuclei (Preformed nuclei + PD166285) conditions. Black dashed lines show the start of each cycle; transparent dashed lines show peaking tubulin intensity, indicating cytoplasmic oscillations when the nuclei are not formed.
